# Therapeutic Targeting of the TGFβ Pathway Augments CAR T cell Efficacy in Glioblastoma

**DOI:** 10.64898/2026.10.25.759450

**Authors:** Darya Alizadeh, Robyn A. Wong, Madeleine S. Maker, Dominic J. Bellavari, Jonathan C. Hibbard, Heini M. Natri, Wuyang Chen, Min-Hsuan Chen, Xiwei Wu, Yate-Ching Yuan, Joseph R. Pecoraro, John D. Jeppson, Renate Starr, Zhiqiang Wang, Nicholas E. Banovich, Stephen J. Forman, Behnam Badie, Christine E. Brown

## Abstract

Chimeric Antigen Receptor (CAR) T therapy represents a promising approach to improve outcomes for patients with high-grade gliomas (HGG). Recently, we reported on a phase I trial assessing the local delivery of CAR T cells for HGG. Despite notable clinical benefits to CAR T treatment, clinical responses were highly variable between patients. Here we report that TGFβ levels in the tumor cavity fluid (TCF) decreased following CAR T treatment and that lower TGFβ levels positively correlated with improved patient responses to CAR T therapy. Patients were stratified based on baseline TGFβ levels in the TCF before CAR T therapy. Single-cell RNA sequencing of patient tumor tissues identified significant upregulation of genes associated with angiogenesis and epithelial-to-mesenchymal transition, and IL6 and SLRP pathways in TGFβ^high^ patients, while IL2, chemokine signaling and NK-mediated cytotoxicity pathways were significantly upregulated in TGFβ^low^ patients. Preclinically, we demonstrated that systemic treatment with a TGFβR1 inhibitor markedly improved CAR T efficacy in syngeneic murine glioma models, correlating with changes in the tumor microenvironment (TME) that benefited both endogenous and CAR T cell phenotype and trafficking. Further studies demonstrated that inhibition of the TGFβ pathway in CAR T cells, through either TGFβR1 inhibitor or CRISPR/Cas9 knockout of TGFβR2 (TGFβR2-KO), significantly enhanced CAR T cell expansion and antitumor activity. Collectively, our clinical and preclinical findings strongly indicate that targeting the TGFβ pathway could significantly enhance CAR T therapy for patients with high-grade gliomas.

**One Sentence Summary:** High TGFβ levels in tumor is linked to resistance to CAR T therapy in GBM patients supporting targeting this pathway to enhance CAR T therapy for GBM.

## INTRODUCTION

Despite promising outcomes from chimeric antigen receptor (CAR) T cells in hematological malignancies, several challenges limit the success of CAR T therapy in solid tumors. In glioblastoma (GBM), CAR T therapy has demonstrated feasibility and safety with signs of efficacy in selected patients in early phase I clinical trials(*1, 2*). Our team has also reported feasibility, safety, and promising therapeutic activity of our IL13Rα2-targeting CAR T cells(*3*). While these initial clinical results are encouraging, most clinical responses are transient, and tumor recurrence is often inevitable. The profoundly immunosuppressive tumor microenvironment (TME), intrinsic T cell dysfunction, and tumor antigen heterogeneity have all been implicated as key factors contributing to the limited efficacy of CAR T cell therapy in solid tumors. These limitations necessitate the development of novel strategies to enhance the function, persistence, and specificity of CAR T cells in the context of GBM.

GBM TME, with its suppressive and immunologically cold tumor features (T cell excluded tumors), remains a major barrier and hinders effective immune response. Identifying and targeting factors that regulate T cell exclusion, suppression or dysfunction is critical to rationally identifying strategies to overcome the GBM TME. Among the pathways implicated in driving immunosuppression across solid tumors, including GBM, the transforming growth factor-beta (TGFβ) pathway has emerged as a central player. TGFβ signaling is known to contribute to glioma initiation, progression, invasion, angiogenesis, and the maintenance of glioma stem cell (GSC) populations(*4*). Importantly, it also plays a pivotal role in shaping the immunosuppressive TME by promoting T cell exclusion(*5*), enhancing immune checkpoint expression(*6*), and impairing the function of effector immune cells(*7*). Thus, therapeutic strategies that target the TGFβ pathway may help overcome these barriers and enhance CAR T cell infiltration, function, and persistence within the tumor.

In this study, we aim to enhance CAR T cell therapeutic potency by targeting the TGFβ signaling axis. By mitigating the suppressive effects of TGFβ, we seek to enhance CAR T cell function, persistence, and infiltration, ultimately improving treatment outcomes for patients with GBM. We assessed the blockade of TGFβ signaling, either through blocking the pathway in the glioma TME using TGFβR1 inhibitor treatments or the signaling on CAR T cells via gene knockout of TGFβR2.

## RESULTS

### High TGFβ levels in tumor cavity is associated with poor response to CAR T cell therapy in patients with GBM

One of the barriers to an effective CAR T therapy for solid tumors is the suppressive TME(*8*). The TGFβ pathway is one of the dominant immunosuppressive pathways in solid tumors, including GBM(*4*). Assessing the TCGA database from 155 patients with GBM tumors, we observe upregulation of TGFβ1 gene expression in GBM tumors compared to normal brain tissues (**fig. S1A)**. The expression of pSMAD2 was also significantly upregulated in patient glioma samples relative to normal brain, indicating TGFβ pathway activation in glioma tumors (**fig. S1, B and C**). In line with this observation, gene set enrichment analysis confirms that the TGFβ pathway is highly upregulated in brain cancer tissues relative to normal brain (**fig. S1D**). Conversely, immune-related genes were significantly downregulated in tumor tissues compared to normal brain (**fig. S1E**). Collectively, these observations confirm that the TGFβ pathway is upregulated in gliomas, and that the activation of this pathway is correlated with a concomitant downregulation of immune-related genes.

To assess the clinical impact of TGFβ levels on patient outcomes to CAR T therapy, we queried patient cerebrospinal fluid (CSF) and tumor cavity fluid (TCF) samples from our recently completed phase I trial evaluating IL13Rα2-targeted CAR T cells in recurrent high-grade glioma (NCT02208362). In this trial, 65 patients were treated with CAR T cells administered locoregionally via surgically-placed catheters through three delivery routes: 1) intratumorally (ICT), 2) intracerebroventricular (ICV), or 3) dual (both ICT and ICV), which enabled access to both TCF and CSF for correlative assessments (**Fig. 1A**). Liquid biopsy of the CSF and TCF, prior to and during CAR T therapy when clinically feasible, enabled measurements of cytokines, including TGFβ. We observed low TGFβ levels in the CSF ranging from 0.5 - 600 pg/mL (log10conc = -0.3 - 2.76; median 31.6 pg/mL), and higher levels measured in the TCF ranging from 12-13120 pg/ml (log10conc = 1.1 – 4.11, median = 754 pg/mL) (**Fig. 1B**). Evaluating the change in the TGFβ levels post CAR T cell therapy, we found that TGFβ levels in TCF were significantly reduced post-CAR T therapy (average decrease 325 pg/mL) while no significant decrease was observed for TGFβ levels in CSF (**Fig. 1C**). Focusing on patients (n=17) with available samples to measure TGFβ levels in the TCF prior to CAR T therapy, we identified an association between TGFβ levels and patient clinical response to CAR T therapy, with progressive disease patients having the highest levels of TGFβ in the TME (**Fig. 1D**). The dynamic changes in TGFβ levels pre-CAR compared to early (1 day after) or late (>2days after) post-CAR T therapy indicates that the impact of CAR T therapy on TGFβ levels is not transient and there is a prolonged effect on the TGFβ level changes in this patient cohort (**Fig. 1E**).

**Fig. 1.**
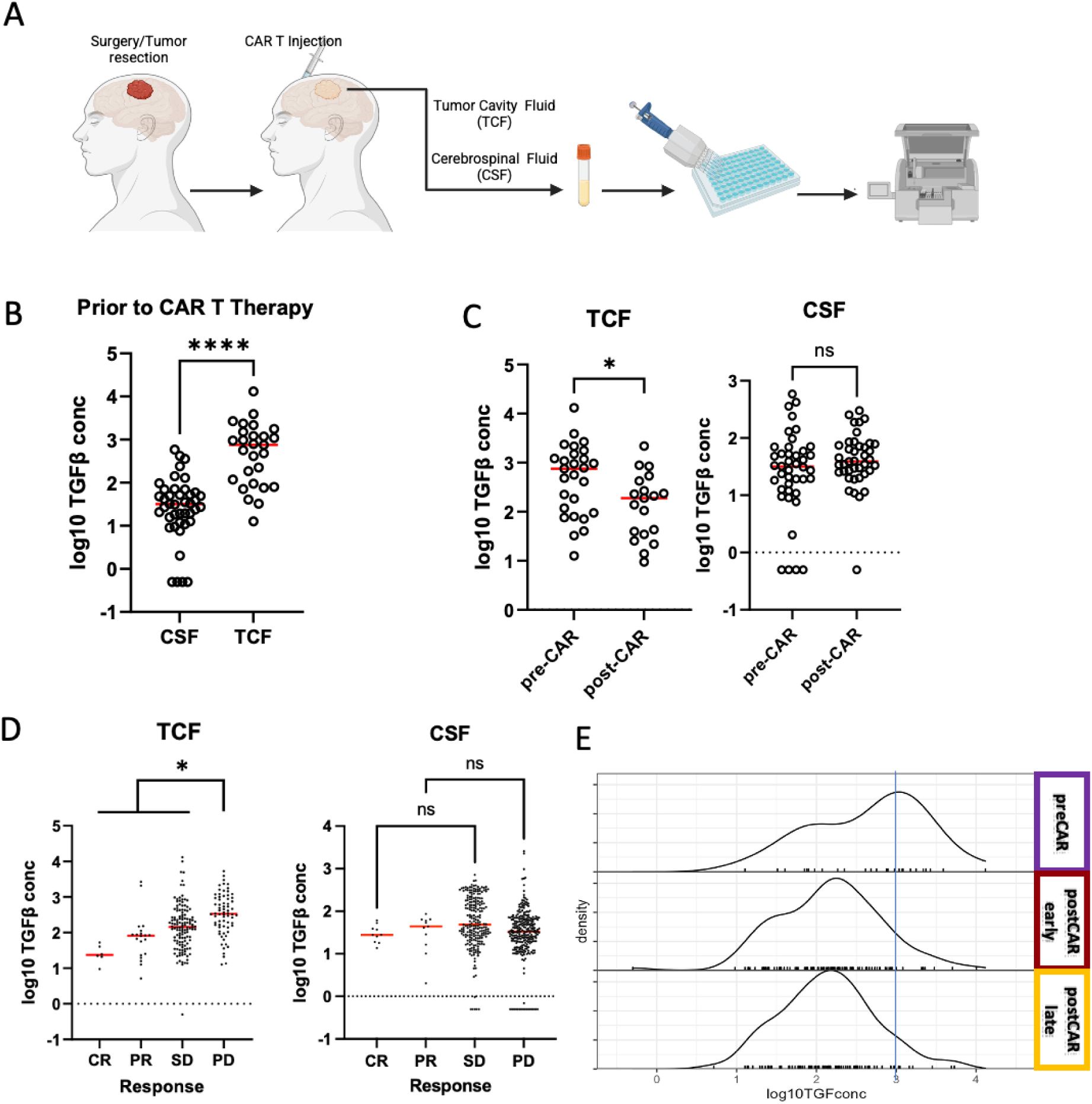
TGFβ levels in patients treated with CAR T therapy. (**A**) Schematic of the samples collection. Samples from tumor cavity fluid (TCF) and cerebrospinal fluid (CSF) are collected post-surgery, pre- and post-CAR T treatment at each cycle. (**B**) TGFβ levels in TCF and CSF of patients with GBM prior to CAR T therapy. (**C**) Comparison of TGFβ levels in TCF (left) or CSF (right) before and after CAR T treatment. (**D**) Correlation between TGFβ levels in TCF (left) and CSF (right) in relation to clinical response to CAR T therapy. (**E**) Density plot shows dynamic changes in TGFβ levels in TCF at during the treatment cycle (preCAR = C1D0; post CAR early = CxD1 or CxD2; post CAR late = CxD3+) CR: complete response; PR: partial response; SD: stable disease; PD: progressive disease.

### TGFβ levels in GBM patients correlate with distinct TME and CAR T cell profile

High expression of TGFβ is associated with immunosuppressive, T cell excluded TME and resistance to immunotherapies, such as checkpoint blockades(*5, 9*). TGFβ levels in the CSF in humans without major CNS disease has been reported to be ranging from ∼30–140 pg/mL(*10*). Accordingly, patients with available samples and TGFβ measurements in the TCF (n=17) were stratified based on their TGFβ levels at baseline prior to CAR T therapy (TGFβ ^low^∼ < 200 pg/mL; TGFβ ^high^= > 400 pg/mL). To assess whether there are any unique features in the TME of patients with differing TGFβ level in the TCF, scRNAseq from tumor tissues of patients (n=17) were also evaluated for pathway changes (**Fig. 2A**). We obtained analyses conducted by our team(*11*) which identified tumor, myeloid and lymphocyte populations. In the lymphoid compartment of TGFβ ^low^ patient cohort, we identified upregulation (p < 0.001) of genes associated with T and B cells (*CD3G, CD8A, CD19*), activation and exhaustion (*CD38, STAT1, IL21, TOX, CD28, ICOS and TIGIT*) and resident migratory T cells (*CXCR3, CXCR6, and ITGAE*), which was concomitant with pathways upregulated in IL2, chemokine signaling and NK-mediated cytotoxicity in the lymphoid population, a signature characteristic of a microenvironment primed for a robust immune response (**Fig. 2B**). In contrast, patients with TGFβ^high^ had upregulation of genes such as TGFβ, *CXCR4, CXCL1, CCL2*, and CCL20 in the lymphoid subsets. Accordingly, in the myeloid compartment, an upregulation of genes associated with myeloid activation and antigen presentation (i.e. *HLA-DRA, CD74, TAP1, CD40, CD86, and CXCL9*) was detected in TGFβ^low^ patient cohort which was associated with significant upregulation of IFNα, IFNγ, and antigen processing and presentation pathways in the myeloid cells. In the TGFβ^high^ cohort, however, myeloid population exhibited significant upregulation of genes involved in suppressive phenotype (i.e. *CXCL1, MRC1, CXCL2, S100A9.A4 and CXCL3*) (**Fig. 2C**). Pathway enrichment analyses in both myeloid and lymphoid populations revealed significant enrichment for genes associated with angiogenesis and epithelial-to-mesenchymal transition in TGFβ^high^ patient cohort, both of which are characteristic of more aggressive tumors and poor prognosis (**Fig. 2, B and C**). These findings identify a common gene signature and pathway enrichment among the immune cell populations associated with high or low TGFβ levels in glioma TME and prognosis.

**Fig. 2.**
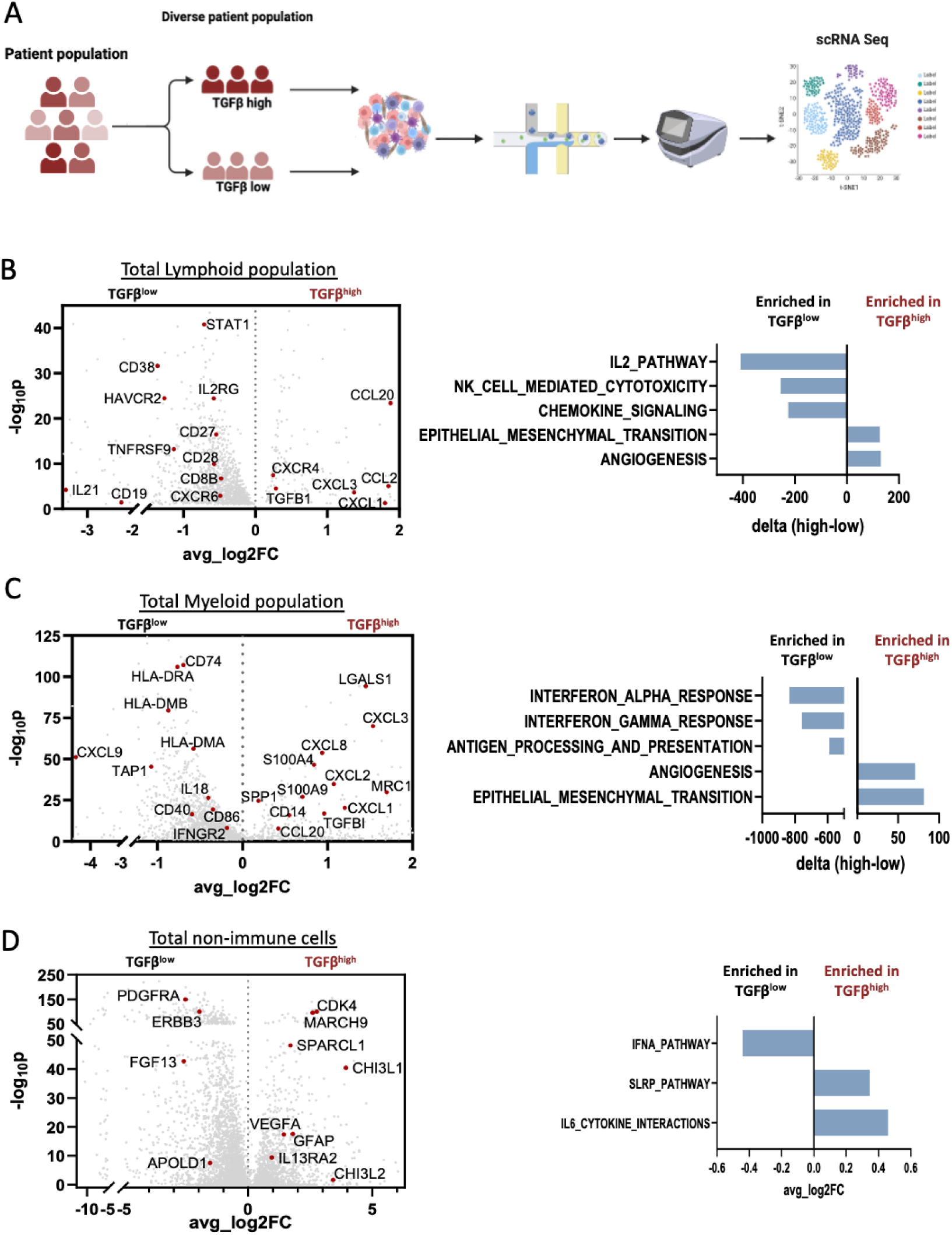
scRNAseq reveals differences in patient tumor microenvironment with different TGFβ levels. Schema showing stratification of TGFβ levels followed by scRNAseq of patient tumors before CAR T treatment. The lymphoid (**B**) myeloid (**C**), and non-immune (**D**) cell compartments were analyzed separately. Volcano plots (left) showing differential expression of genes in TGFβ^high^ or TGFβ^low^ cohorts. Bar graphs (right) show differentially expressed pathways from gene set enrichment analysis.

In the non-immune populations (tumor and fibroblast), we identified genes associated with extracellular matrix (ECM) changes and mesenchymal phenotype, such as *CHI3L1*, *CHI3L2*, *VEGF*, and *IL13RA2* genes significantly upregulated in the TGFβ^high^ group. Furthermore, pathway enrichment analysis showed significant upregulation of IL6 and SLRP pathways, in non-immune cell population of TGFβ^high^ patients suggesting a correlation between high levels of TGFβ and pathways involved in chronic inflammation, angiogenesis, proliferation and migration/invasion of gliomas while the TGFβ^low^ cohort had significant enrichment in type I IFN signaling (**Fig. 2D**).

### Blockade of TGFβ pathway in CAR T cells improves CAR T phenotype and function

Based on our clinical findings, we sought to evaluate the effect of targeting the TGFβ pathway in CAR T cells. We first showed that blocking the TGFβ pathway with TGFβR1 inhibitor (TGFβR1i [LY3200882]) (**Fig. 3, A and B**) significantly restored TGFβ-mediated suppression of human T cell proliferation (2-fold) (**Fig. 3C**) and effector phenotype (expression of genes i.e. *PRF1*, *GZMA,* and *IFNγ*) and decreased TGFβ-mediated upregulation of PD1 (**Fig. 3, D to F**). Furthermore, we performed a recursive killing assay in which CAR T cells were cocultured with IL13Rα2-expressing tumor cells and subsequently rechallenged with additional tumor every 2 days for 7 days(*12*). We demonstrated that while TGFβ decreased CAR T expansion and function, blockade of TGFβ signaling significantly restored CAR T expansion and antitumor function in both high and low antigen tumor models (**Fig. 4, A and B; fig S2)**. To characterize the underlying phenotypic changes, CAR T cells were isolated after recursive killing and CD4/CD8 populations were separated for RNA sequencing using Nanostring technology. Gene expression analysis showed that TGFβ exposure downregulated genes associated with T cell activation and cytotoxic function (e.g., *GZMA*, *PRF*, *CD40L*, *CD127*, *CD28*, etc.), while blocking TGFβ signaling with TGFβR1i restored the expression of these genes. Additionally, TGFβ-mediated upregulation of genes linked to regulatory and inhibitory functions (e.g., *IL4R*, *CLTA4*, *TIGIT*, *IKZF2*, *NR3C1*, *and CCL22/CCR4*) was significantly reduced upon TGFβR1i treatment. The phenotypic changes were observed in both CD4 and CD8 subsets (**Fig. 4C).** Furthermore, in T cells treated with the TGFβR1 inhibitor, pathway enrichment analyses revealed an upregulation of genes associated with memory CD8 and naïve CD4 T cells, and downregulation of T cell exhaustion and Th2 gene signature (**Fig. 4D**) implying that inhibition of TGFβ signaling promoted memory T cell phenotype and reduced T cell exhaustion and Th2 type CD4 cells. Together these results support the notion that blocking TGFβ signaling improves phenotype, enhances expansion and antitumor function of CAR T cells, thereby further rationalizing a promising therapeutic approach for future of CAR T therapy for GBM.

**Fig. 3.**
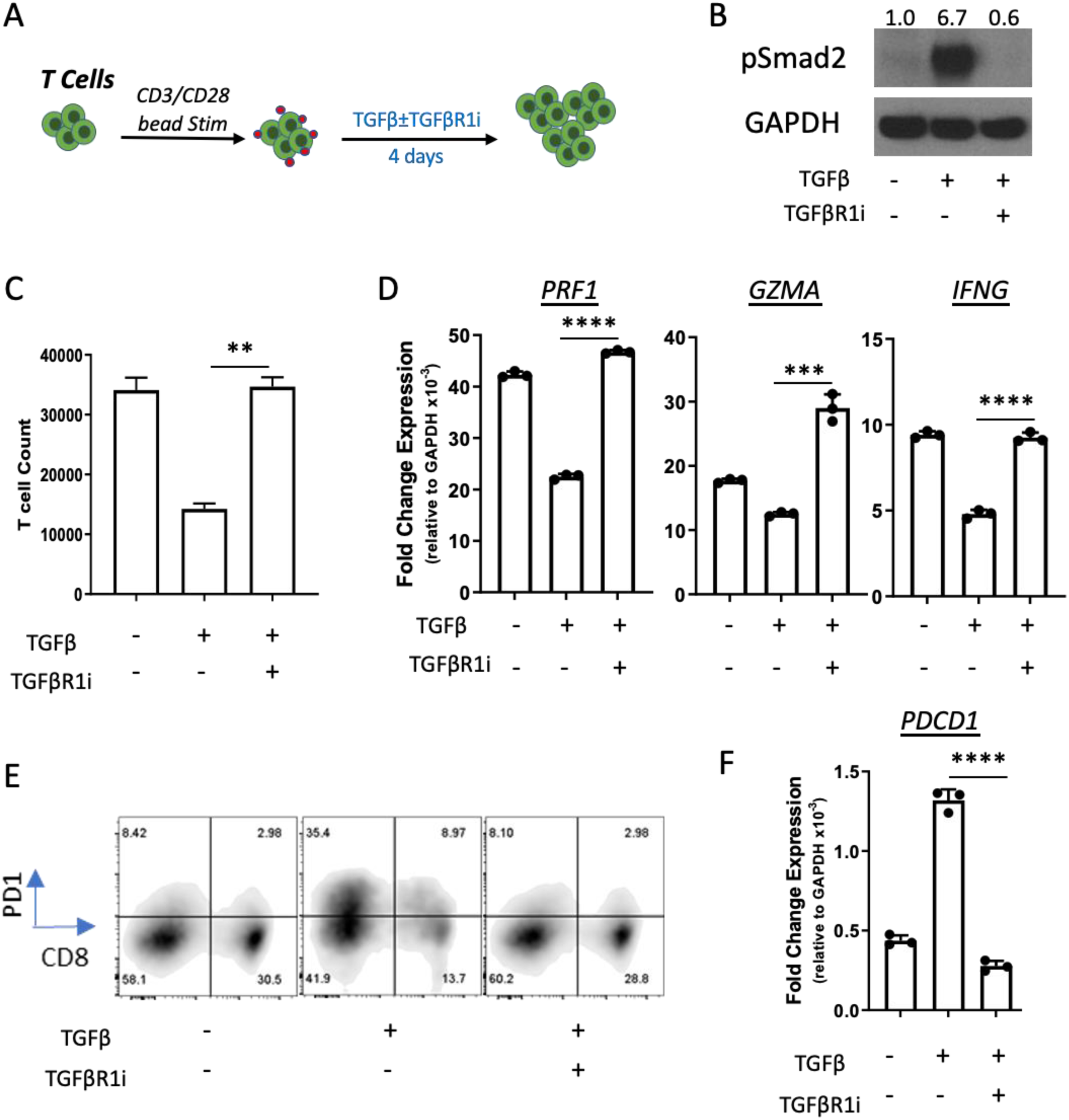
Blocking TGFβ signaling in activated T cells. (**A**) Schematic of experimental design. (**B**) Western blot analysis shows levels of pSmad2 (S465/467) and GAPDH proteins. Numbers indicate ratio of pSmad2 to GAPDH by densitometry, normalized to the non-stimulated control. (**C**) Bar graph shows T cell proliferation upon CD3/CD28 bead stimulation with or without TGFβ (10 ng/ml) and TGFβR1 inhibitor (3 µM). (**D**) Graphs show expression of genes associated with cytotoxic function (perforin:*PRF1*; granzyme A: *GZMA*; interferon gamma: *IFNG*). (**E**) Flow cytometry demonstrates expression of PD1 protein and (**F**) Bar graph shows expression of PD1 gene (*PDCD1*) in bead stimulated T cells with or without TGFβ (10 ng/ml) and TGFβR1 inhibitor (3 µM) after 4 days.

**Fig. 4.**
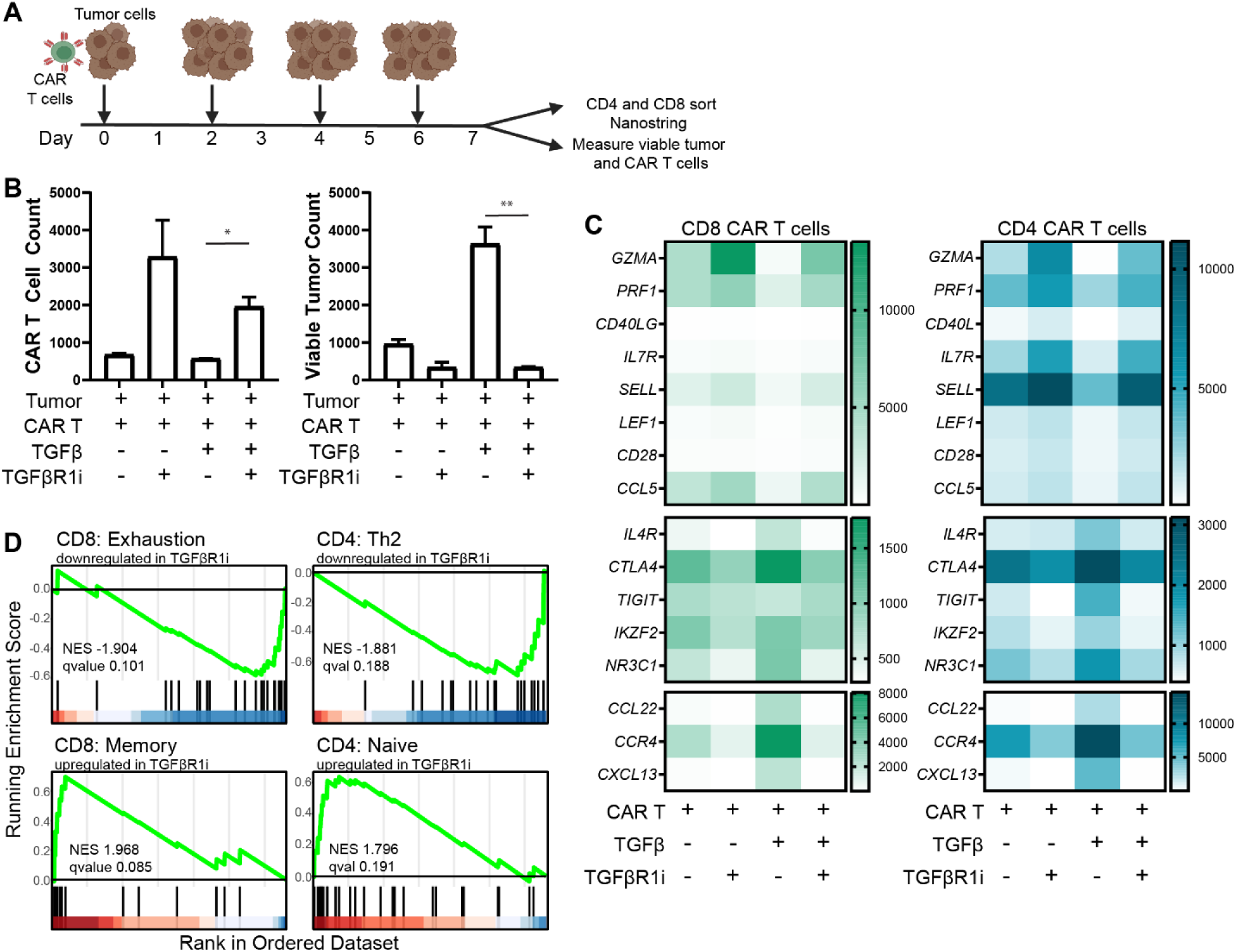
Blocking TGFβ signaling enhances CAR T expansion and function. (A) Schematic of experimental design. (**B**) Graphs show CAR T cell expansion (left) and viable tumor counts (right) after multiple tumor rechallenge in the presence or absence of TGFβ (10 ng/ml) or TGFβR1 inhibitor (3 μM), after coculture for 7d at an initial 1:4 E:T ratio, against a patient derived glioma tumor expressing high levels of IL13Rα2. (**C**) Heatmaps show key gene expression differences as determined by NanoString platform, in remaining live T cells, which were sorted by flow cytometry into CD4 and CD8 populations. (**D**) Enrichment plots show differences in T cell phenotype in either CD4 or CD8 populations with TGFβR1 inhibitor (3 µM, with 10 ng/mL TGFβ) over TGFβ (10 ng/mL) alone.

### Blockade of TGFβ signaling enhances CAR T therapy in syngeneic immunocompetent **glioma tumors**

We next sought to understand the importance of blocking TGFβ pathway in the TME. To determine the impact of blocking TGFβ signaling in the TME, we used our well-established, immunocompetent syngeneic glioma model(*13*). Murine KR158-IL13Rα2+ glioma tumors (K-Luc) were allowed to fully establish for 14 days post-implantation, a time point at which CAR T cell treatment typically provides only a modest therapeutic effect (**fig. S3**). Tumor bearing mice were treated with TGFβR1 inhibitor (TGFβR1i; LY3200882), CAR T cells, or a combination of TGFβR1i with CAR T therapy (**Fig. 5A**). Both CAR T cells and TGFβR1i as monotherapies had modest effect and failed to significantly improve the overall survival of the tumor bearing mice (**Fig. 5, B to D)**. However, the combination of TGFβR1 inhibitor followed by CAR T therapy resulted in significant regression of the tumor and enhanced overall survival by 65-70% (**Fig. 5, B to D)**. Importantly, there were no signs of toxicity or treatment-associated weight loss in any groups (**fig. S4**). Similar findings were observed in a second syngeneic glioma model (GL261 IL13Rα2^+^) (**fig. S5, A and B**), further highlighting the TGFβ pathway as a dominant pathway in GBM and confirming the importance of targeting this pathway for CAR T cell therapy. Surviving mice that were tumor-free were rechallenged with antigen negative parental tumors showed 100% survival (**fig. S5, A to C)** (*13*). To elucidate immune-related changes in the TME that coincided with the rejection of antigen negative tumor, brain samples post tumor rechallenge were harvested and analyzed. Assessment of TME after challenge with antigen negative tumors revealed ∼2-fold increase frequency and number of CD3+ (CD19-) endogenous T cells that were activated (CD3+ CD69+) and specifically more CD8 T cells (CD8+CD69+) than CD4 (**fig. S6C**). Gene expression analysis of the TME revealed increase expression of genes associated with activation, cytotoxic and effector function (*IFNγ*, *GZMA, CD74,* and *CD86*) (**fig. S6D**). The increase in endogenous T cell counts, specifically activated CD8 T cells, which coincided with rejection of antigen negative tumors strongly indicates the induction of CD8 T cell memory response post combination therapy.

**Fig. 5.**
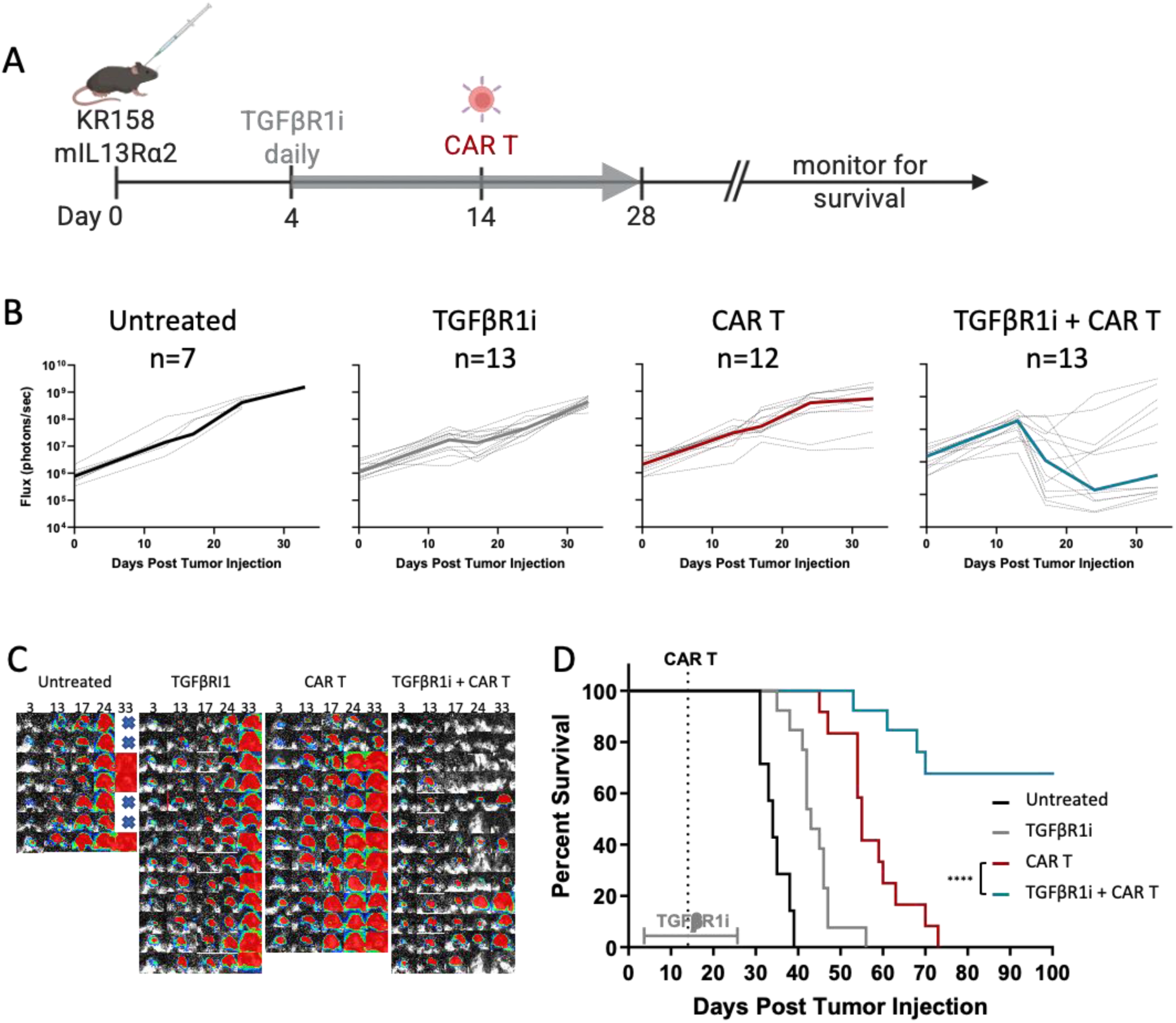
Blockade of TGFβ signaling in the TME improves efficacy of CAR T therapy in a large established glioma model. (**A**) Schematic of experimental design. Mice bearing glioma tumors (14-day old KR158 mIL13Rα2+) were treated with TGFβR1 inhibitor daily before and after IL13Rα2-CAR T treatment. (**B**) Flux graph shows tumor growth kinetics demonstrate changes in tumor progression over time. (**C**) Bioluminescent images show tumor growth kinetic over time. (**D**) Kaplan-Meier shows overall survival of mice receiving monotherapy (TGFβR1 inhibitor alone or CAR T alone) compared to combination therapy (TGFβR1 inhibitor plus CAR T therapy). Data are presented as means ± SEM and were analyzed by two-tailed, unpaired student’s t test. *, *p* < 0.05; **, *p* < 0.01; ***, *p* < 0.001 for indicated comparison.

#### Inhibition of TGFβ signaling changes the glioma tumor immune microenvironment

Presence of TGFβ in the tumor has been associated with suppression of effector T and NK function and development of an immune-excluded TME(*5*),(*14*). To understand the changes in the TME that corresponds to the synergistic effects of combining TGFβR1i with CAR T therapy and improved survival, tumor tissues were analyzed 3 days after treatment (**Fig. 6A**). To characterize T cell populations and differentiate changes in endogenous versus adoptively transferred T cells at cellular level, isogenically mismatched CD45.1 CAR T cells were used(*13*). While the combination therapy showed higher CAR T cell count (CD3+ CD45.1+CD19+) compared to CAR T alone, both treatment groups exhibited comparable activation levels (CD45.1+CD19+CD69+), suggesting antigen-dependent CAR T activation (**Fig. 6B**). Endogenous T cell count (CD3+CD45.1-) was not significantly different at early timepoint (3 days); however, there was a 40% increase in frequency of activated (CD69+) endogenous T cells in combination CAR T plus TGFβR1i group as compared to each monotherapy or untreated groups **(Fig. 6C**). Since TGFβ has been linked to T cell exclusion in the TME(*5, 14*), we next assessed the expression of CXCR3+, a chemokine receptor involved in trafficking and migration. Interestingly, the combination TGFβR1i plus CAR T treatment showed an increase in frequency of CXCR3+ T cells in both CAR and endogenous T cells (**Fig. 6D**). Mice receiving TGFβR1i alone also demonstrated 20% increase in CXCR3+ endogenous T cells, albeit not statistically significant, indicating that TGFβ inhibition induced expression of CXCR3+ in T cells which indicates improve T cell trafficking (**Fig. 6D**). In addition, inhibiting TGFβ pathway increased the frequency of cytotoxic and proliferating T cells (CD3+IFNγ+Ki67+) compared to CAR T or TGFβR1i alone (**Fig. 6E and fig. S7**).

**Fig. 6.**
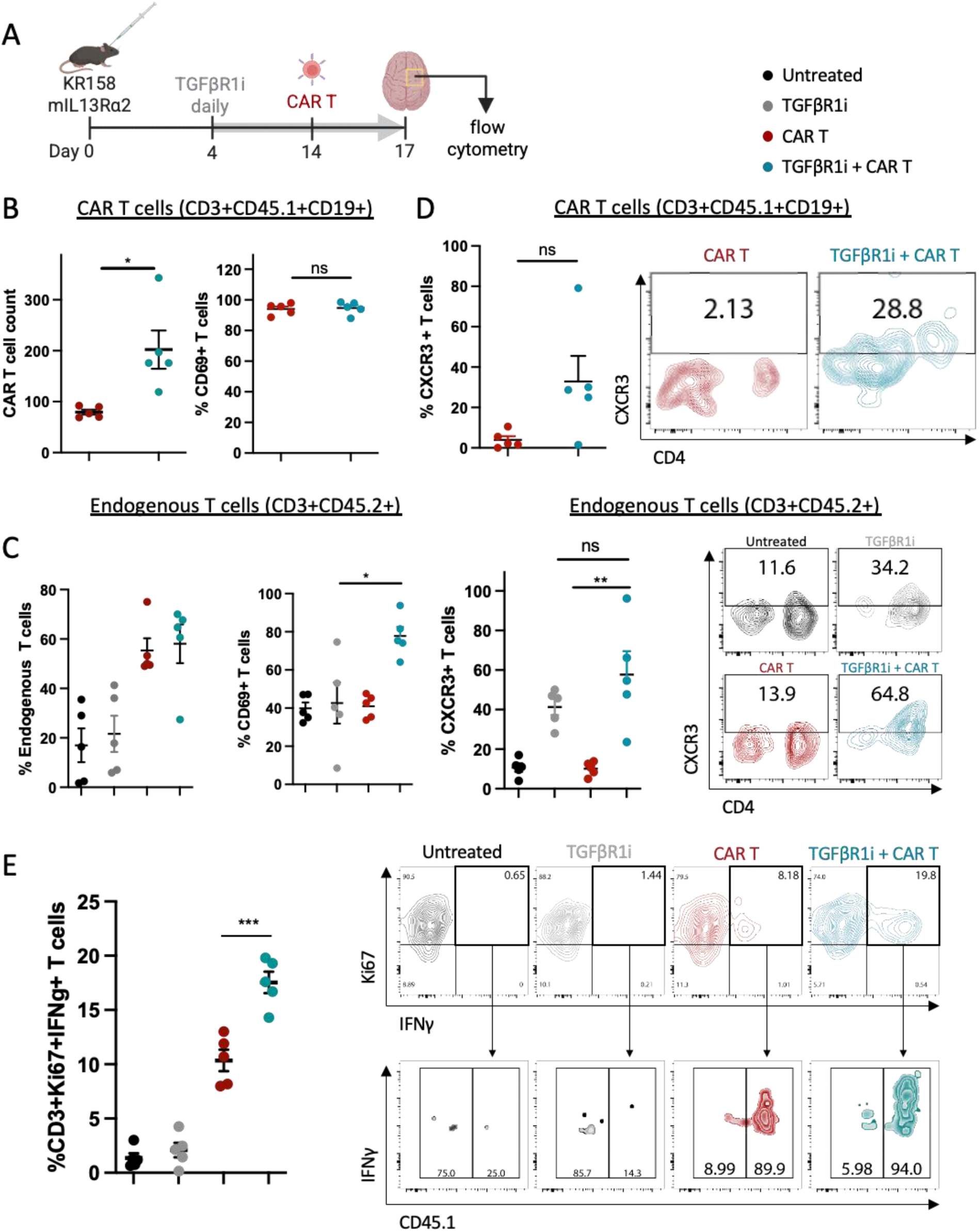
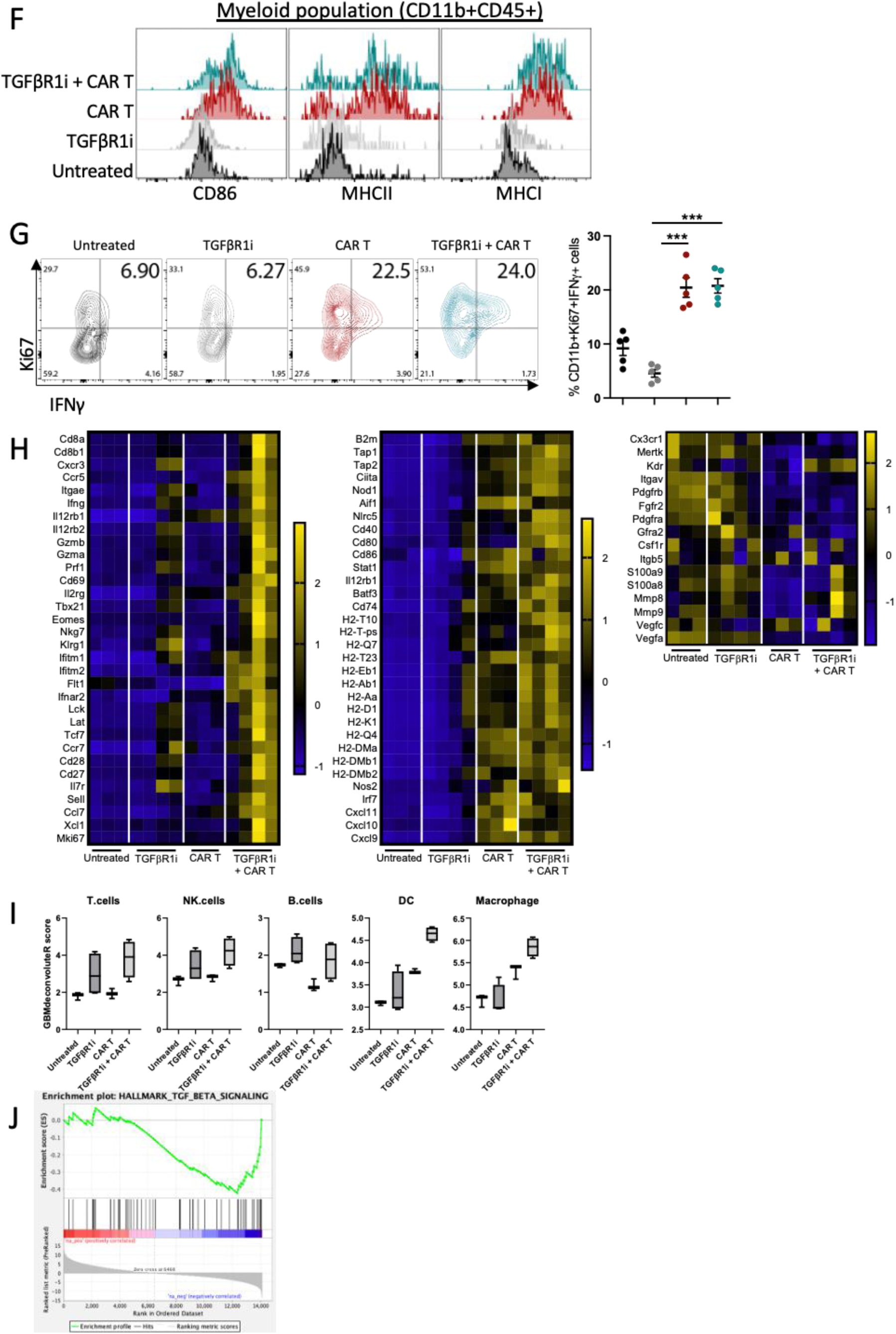
Inhibition of TGFβ signaling in combination with CAR T therapy changes the tumor microenvironment. (**A**) Schematic of experimental design. (**B**) Graphs quantify flow cytometry analysis of CAR T cell phenotype from CAR T treated or combination group. (**C**) Graphs quantify flow cytometry analysis of endogenous T cell and representative dot plots. (**D**) Graphs quantify flow cytometry analysis of CXCR3 expression on CAR T cells (top left) or endogenous T cells (bottom left). Contour plots show representative flow cytometry from CAR T treated (top right) or endogenous T cells (bottom right). (**E**) Graphs quantify flow cytometry analysis of IFNγ and Ki67 on total T cells from the tumor microenvironment (left) with representative flow cytometry plots (right). (**F**) Histograms of mean fluorescence intensity of the indicated markers on the myeloid population, measured by flow cytometry in each treatment groups. (**G**) Flow cytometry analysis of myeloid cells from each treatment group. (**H**) Heatmaps show mRNA expression of tumor microenvironment for each treatment group. (**I**) Deconvolution of RNA seq from tumor identifies different immune populations in each treatment group. (**J**) Pathway enrichment analysis shows upregulation of TGFB pathway in tumor. Data are presented as means ± SEM and were analyzed by two-tailed, unpaired student’s t test. *, *p* < 0.05; **, *p* < 0.01; ***, *p* < 0.001; ****, *p* < 0.0001 for the indicated comparison.

Evaluating changes in resident microglia/macrophage populations by flow cytometry revealed an increased frequency of activated brain-resident macrophage/microglia cells (CD86+, MHCI^high^, MHCII^high^) in groups treated with CAR T cells (either CAR T alone or combination CAR T plus TGFβR1 inhibitor) (**Fig. 6F**). Similarly, induction of effector myeloid cells (CD11b+ IFNγ+Ki67+) was observed only in CAR T and combination therapy groups (**Fig. 6G**). The activation of myeloid compartment with CAR T cell treatment, irrespective of TGFBR1i treatment, confirmed our previous findings of CAR T-mediated local activation of resident myeloid cells in the brain. Of note, no significant impact or contribution of TGFβR1i to the myeloid cell phenotype was observed.

Furthermore, to increase resolution and evaluate changes in the TME, we performed RNA sequencing analysis of the bulk tumor tissue 3 days after treatment. Evaluating lymphoid related gene expression, the group that received the combination therapy exhibited a significant upregulation of genes associated with T cell activation and effector function (*Gzma, Gzmb, Eomes, Tbx21, Cd40, Ifng, Cd69*) and genes associated with T cell trafficking (*Cxcr3*, *Ccr5*) and memory phenotype (*Cd27*, *Tcf7*, *Ccr7*) (**Fig. 6H**). We previously showed that local treatment of CAR T cells activates resident myeloid populations(*13*), indeed here we also observe a significant induction of genes associated with macrophage activation, antigen presentation and effector function (*Cd80, Cd86, Cd74, Cxcl9/10, Nos2* (iNOS)) in groups that received CAR T treatment (CAR T alone or combination TGFβR1i plus CAR T) (**Fig. 6H**). In contrast resident myeloid cells from untreated and TGFβR1i treated groups exhibited high expression of *Mrc1 (*CD206*), S100a8/9, Csfr1, Vegfa*—markers frequently associated with anti-inflammatory, immunosuppressive macrophages (**Fig. 6H**). Deconvolution of the TME revealed higher frequency of genes associated with lymphocyte populations, T, NK and B cells, in groups that received TGFβR1i, and genes associated with myeloid populations, macrophages and dendritic cells (DC), in groups receiving the CAR T and combination therapy emphasizing the importance of combining TGFβ blockade with CAR T cells (**Fig. 6I**). Pathway enrichment analysis confirmed that genes associated with TGFβ signaling were downregulated in combination therapy (**Fig. 6J**). These findings show that CAR T cells and TGFβR1i each contribute to the immune landscape in distinct ways--CAR T cells driving activation of myeloid cells and changing their phenotype while TGFβR1i enhanced T cell activation and migratory phenotype. Collectively, these results confirm that CAR T therapy and TGFβR1i treatment remodel the TME to exhibit features characteristic of antitumor immune responses.

### Knocking out TGFβ pathway in CAR T cells enhances CAR T antitumor function

Based on our findings that inhibiting TGFβ signaling with TGFβR1 inhibitor significantly altered CAR T cell phenotypic characteristics, expansion, and antitumor activity, we sought to block TGFβ signaling in CAR T cell products by genetically knocking out TGFβR2. We generated TGFβ resistant CAR T cells by knocking out TGFβR2 (TGFβR2KO) in murine CAR T cells using CRISPR/Cas9 (**Fig. 7A and fig. S8**). TGFβR2KO and wild-type (wt) CAR T cells exhibited comparable transduction efficiency, viability, and expansion (**fig. S8**). *In vitro*, TGFβR2KO-CAR T cells demonstrated superior antitumor activity (50% more antitumor function), and a higher T cell count (40% more expansion) compared to wild-type (wt) CAR T cells (**Fig. 7B**). To determine if blocking TGFβ signaling in CAR T cells improves their function, we used the syngeneic glioma models described previously (*13*) (**fig. S3**). Fourteen days post tumor implantation, murine IL13Rα2-CAR T or TGFβR2KO-IL13Rα2-CAR T were locally (intracranially [IC]) delivered (**Fig. 7C**). Bioluminescent (BLI) images pre and post treatment showed reduction in tumor size in the group that received TGFβR2KO-IL13Rα2-CAR T cells (**Fig. 7D**). Kaplan-Meier survival curve showed a significant delay in survival of mice that received TGFβR2KO-IL13Rα2-CAR T cell (**Fig. 7E**), indicating superior function. Importantly, no signs of toxicity or treatment related weight loss was observed (**Fig. 7F**). Together, these results show a benefit in blocking the TGFβ pathway in CAR T cell design and an opportunity to translate next generation TGFβ resistant CAR T cells (TGFβR2KO-CAR T) to the clinic (NCT06815029).

**Fig. 7.**
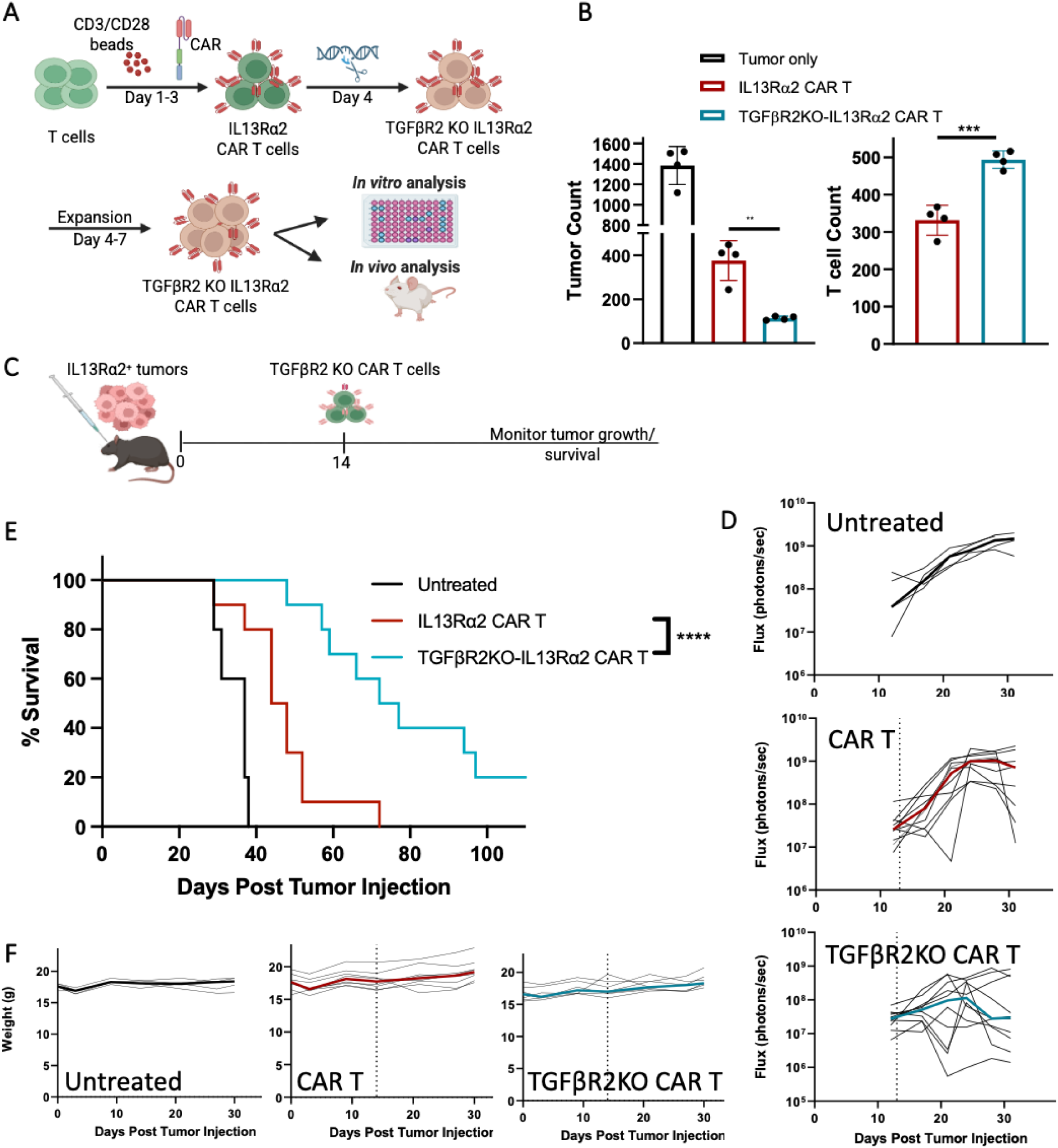
TGFβR2KO-IL13Rα2 T cells antitumor function *in vitro* and *in vivo* in syngeneic GBM model. (**A**) Schematic of *in vitro* manufacturing of CRISPR/Cas9 knockout of TGFβR2 in IL13Rα2-CAR T cells (TGFβR2KO-IL13Rα2-CAR T cells). (**B**) Graph shows tumor count (left) and CAR T expansion (right) after coculture with different IL13Rα2-CAR T cell variants at 1:4 E:T ratio. (**C**) Schema of *in vivo* experimental design. Mice bearing large established tumors (14-day engraftment) were treated with TGFβR2KO or standard-IL13Rα2-CAR T cells. (**D**) Flux graph shows tumor growth kinetics over time. (**E**) Kaplan-Meier survival curve of mice treated with different IL13Rα2-CAR T cell variants. (**F**) Graphs show changes in body weight measured over time.

We next set out to understand the relative contribution of TGFβ-resistance engineered as part of the CAR T cell product versus more global TGFB-inhibition through small molecule pathway inhibition. While TGFβR2-KO prolonged survival as compared to CAR T cell control group, the including of TGFβ pathway blockade with LY3200882, significantly enhanced the survival of both WT CAR T cells and TGFβR2-KO CAR T cells. This indicates that while blocking TGFβ pathway in CAR T cells is beneficial, the inhibition of the pathway in the glioma TME can further enhance glioma antitumor responses (**Fig. 8**).

**Fig. 8.**
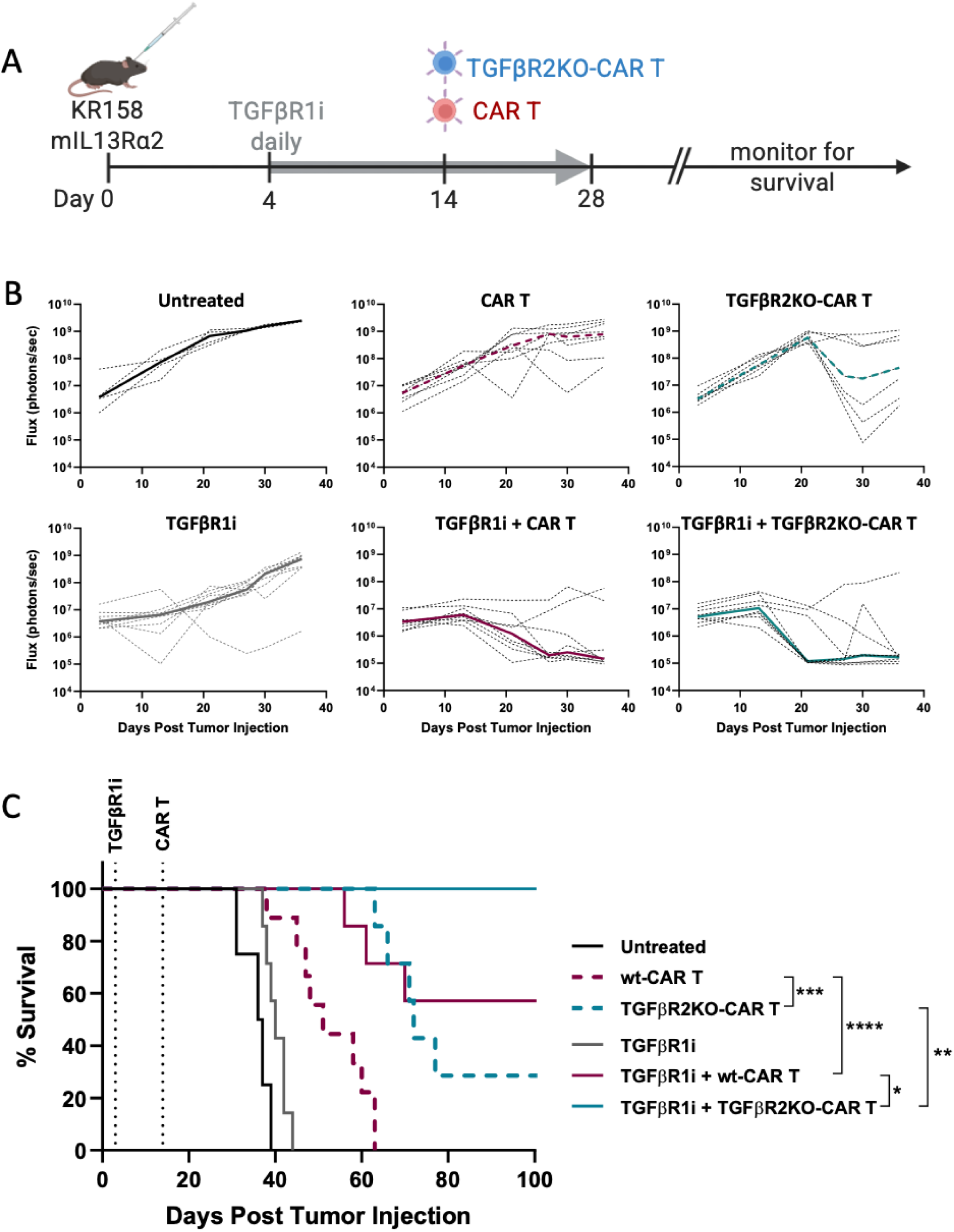
Inhibition of TGFβ pathway in CAR T and TME enhances antitumor efficacy. (**A**) Schematic of experimental design. (**B**) Flux graphs show tumor growth kinetics over time. (**C**) Kaplan-Meier survival curve of mice treated with different IL13Rα2-CAR T cell variants.

## DISCUSSION

Effective CAR T cell therapy in GBM will require defining the dominant TME suppressive pathways that limit CAR T cell function and developing rational strategies to overcome them. Studies from our group and others have previously shown that the TGFβ pathway is upregulated in patients with GBM. Consistent with these observations, analysis of patient samples from our recently completed clinical trial(*15*), revealed elevated TGFβ levels in the tumor cavity fluid prior to CAR T treatment, highlighting a suppressive local milieu and supporting TGFβ as a dominant suppressive pathway in GBM. Notably, low baseline TGFβ levels in tumor fluid before treatment were associated with improved clinical response to CAR T therapy. While TGFβ-mediated immune suppression is well established, comparatively little is known about its impact on outcomes for CAR T therapy in GBM.

Increasing evidence suggests that CAR T efficacy in solid tumors is strongly shaped by intrinsic features in the TME, which can exert both anti-tumor and pro-tumor effects(*3, 11*). To define TME features associated with TGFβ, we analyzed scRNA sequencing of patient baseline tumors, stratified by high versus low TGFβ levels in the tumor fluid. Tumors from patients with high TGFβ exhibited immune populations with more suppressed phenotypes and upregulation of angiogenesis and epithelial-to-mesenchymal transition (EMT) programs, consistent with known pleotropic roles of TGFβ in immune suppression and tumor progression. In contrast, tumors from patients with low TGFβ showed increased expression of genes related to immune activation, antigen presentation, and IFN-signaling response pathways, suggesting a more immune-permissive microenvironment. Notably, within the non-immune compartment (tumor cells and fibroblasts), high TGFβ tumors showed increased expression of ECM and glioma mesenchymal subtype-associated genes, including *CHI3L1*/*CHI3L2* and *IL13RA2*, supporting a link between TGFβ-high states, ECM remodeling and a glioma mesenchymal-like phenotype(*16*),(*17*). Collectively, these findings strongly support therapeutic strategies aimed at mitigating the effects of TGFβ and disrupting immunosuppressive network that promotes glioma progression and suppression of immune responses for enhancing CAR T cell therapy. Accordingly, we investigated two approaches to target the TGFβ axis to enhance CAR T therapy for GBM: (1) small molecule inhibition of the TGFβ pathway within the tumor microenvironment (TME) through TGFBR1-inhibition (LY3200882; provided by Eli Lilly and company)(*18*), and (2) engineering CAR T cells to resist TGFβ-mediated suppression through TGFβR2 gene knockout.

Therapeutic targeting of TGFβ signaling has long been pursued as a strategy to remodel the TME, inhibit tumor progression and enhance antitumor immunity. These efforts have led to the development of small-molecule TGFβR1 inhibitors to broadly attenuate TGFβ signaling within the solid tumor TME. Significant among these TGFβR1 inhibitors are TEW-7197 (Vactosertib-MedPacto)(*19*),(*20*), LY2157299 (Galunisertib, Eli Lilly and company)(*21*), LY3200882 (Eli Lilly and company)(*18*), and PF-06952229 (MDV6058-Pfizer)(*22*), compounds which have advanced into clinical trials, and have generally demonstrated safety, feasibility and signals of clinical activity in solid tumors. Therefore, we reasoned that inhibiting TGFβ signaling in TME in combination with CAR T cells would synergize with CAR T therapy by improving both CAR-intrinsic and endogenous antitumor responses. Based on this knowledge, we investigated the use of a small molecule TGFβR1 inhibitor LY3200882 in combination with IL13Rα2-CAR T cell therapy for treatment of GBM. Indeed, we showed an increase in survival for single-agent TGFβR1 inhibition (LY3200882) in murine glioma models and further demonstrated synergy when combined with CAR T cell therapy. The combination therapy remodeled the tumor immune landscape, including increases in the activation and cytotoxic function of T cells (endogenous and CAR). These observations are in line with our previous findings demonstrating the importance of host immune cells in achieving effective CAR cell therapy in solid tumors(*13*).

TGFβ can act as an immune checkpoint regulator, inducing expression of inhibitory receptors like the PD-1/PD-L1 pathway. Reported synergy between TGFβ pathway inhibition and immune checkpoint blockade has provided a rationale for clinical strategies combining TGFβR1 inhibitors with immune checkpoint blockade(*5, 18*). Phase I studies of TGFβ blockade (LY3200882) as monotherapy or combination therapy with chemotherapy or anti-PD-L1 have shown the approach to be safe and well-tolerated, with preliminary evidence of antitumor activity in advanced solid tumors, including gliomas (*18*). In line with these reports, our data also shows that blockade of TGFβ signaling in CAR T cells significantly reduces the expression of genes associated with exhaustion and inhibitory phenotype (*CTLA4, NR3C1, PDCD1, CCR4, and IL4R*) and upregulation of the genes associated with cytotoxic function *(GZMA, PRF1*, *IFNγ*) (*23*). TGFβ-mediated promotion of CAR T exhaustion and inhibition of expansion and cytotoxic function rationalize for the development of CAR T product that is resistant to the negative effects of TGFβ in the TME.

Beyond promoting exhaustion, TGFβ contributes to immune-excluded tumor states that are linked to resistance to checkpoint blockade, radiation, and chemotherapy(*5, 9, 14*). Mechanistically, TGFβ impairs immune cell trafficking through several pathways, including the induction of the PD-1/PD-L1 axis(*5*) and Smad2-mediated suppression of CXCR3, a chemokine receptor critical for immune cell migration and trafficking(*14*),(*5*). TGFβ-mediated downregulation of CXCR3 expression impairs infiltration of immune cells into the tumor. Furthermore, blockade of TGFβ signaling in TGFβR1-deficient CD8+ T cells enhanced their migration to IFN-stimulatory chemokines, including CXCL10(*14*). These findings align with our observation that CAR T cells treated with TGFβR1 inhibitor, LY3200882, exhibited enhanced CXCR3 expression along with gene-expression profiles associated with a more naïve/memory phenotype. Furthermore, *in vivo*, TGFβ pathway inhibition led to upregulation of CXCR3 on both CAR T cells and endogenous T cells within tumor tissues, suggesting enhanced migratory capacity following combination therapy. Because increasing T cell infiltration into the TME is an emerging strategy to overcome the immunosuppressive nature of GBM(*24*), combining CAR T therapy with TGFβ blockade may attenuate immune suppression through complementary mechanisms, including reducing inhibitory/exhaustion programs, inducing myeloid cell activation, and increasing tumor-infiltrating leukocytes. Collectively, these findings underscore the clinical potential of targeting the TGFβ pathway to enhance the efficacy of CAR T cell therapy in brain tumors.

Based on the critical role of TGFβ on suppressing T cell function, next-generation CAR T cell strategies have focused on engineering T cells for TGFβ-resistance(*23, 25–28*). To date, TGFβ resistant CAR T cells have been generated through overexpressing dominant-negative TGFβR (TGFβRDN) or genetic knockout of TGFβR2 (TGFβR2KO). TGFβRDN CAR T cells have shown promise in both preclinical and clinical studies(*25, 26*),(*27, 28*),(*29*). Toxicity due to over-proliferation and excessive expansion of therapeutic product has been reported in patients with advanced prostate cancer treated with TGFβRDN CAR T cells^16^. CRISPR knock out of TGFβR2 has also shown promising therapeutic outcome in preclinical immunocompromised models(*23*). Using our clinically relevant immunocompetent model, we evaluated the efficacy of TGFβ resistant CAR T cells by CRISPR knock out of TGFβR2 in murine CAR T cells. We showed significant enhancement in efficacy in TGFβ-resistant CAR T cells with no signs of treatment-related toxicity providing evidence that in the setting of GBM local regional delivery of TGFβR2KO-CAR T cells is efficacious, and importantly safe and feasible. While developing a TGFβ-resistant CAR T product that can withstand the suppressive TME has great therapeutic potential, our analyses of patient tumors reveled expression of TGFβ in both endogenous lymphoid and myeloid compartment in TGFβ^high^ patient cohort prior to CAR T therapy, underscoring the importance of blocking the pathway in the TME. Indeed, our studies confirm that blocking TGFβ signaling in the TME enhanced both wt CAR and TGFβ KO CAR T cell functions.

Delineating mechanisms that positively modulate antitumor immunity is critical for developing potent CAR T therapies in solid tumors. This work underscores the profound influence of the GBM TME on CAR T efficacy through production of suppressive factors and summarizes two complementary approaches to overcome TGFβ-mediated immune suppression for enhancing CAR T therapy in GBM. We developed two strategies to overcome TGFβ suppressive impact by engineering into the T cell themselves: “resistance to suppression” or by including pharmacological intervention to block the pathway and promote CAR and/or endogenous T cell antitumor activity. The prospect of enhancing CAR T therapy by blocking TGFβ signaling in patients with GBM shares similar immunosuppressive pathways as other solid tumors; therefore, targeting this pathway may be a promising approach for treatment of a broad spectrum of other solid tumors with highly suppressive TME. Our findings have provided a solid foundation for clinical translation of incorporating TGFβ blockade during CAR T therapy, which as a result, both strategies (blockade at CAR or TME) are at different stages of clinical application for patients with recurrent GBM. We are clinically exploring the utility of TGFβ-blockade on CAR therapy in GBM and have initiated a phase I clinical trial evaluating TGFβR2-KO CAR T cells and are currently enrolling patients with recurrent GBM (NCT06815029). As our data also highlights GBM TME characterized by elevated TGFβ levels, which significantly limits CAR T therapy, strongly rationalizing the incorporation of small molecule TGFβ pathway inhibitor in combination with CAR T therapy. While Eli Lilly and company is no longer pursuing clinical application of LY3200882, we are currently pursuing other clinically tested small molecule TGFβ pathway inhibitors that have shown safety and efficacy with goal of initiation a clinical trial in combination with CAR T therapy in the near future. Data obtained and lessons learned from these trials will inform not only future trial designs for GBM patients but other solid tumors that have limited therapeutic options.

## MATERIALS AND METHODS

### Mice and Cell lines

C57BL/6/J or CD45.1 (B6.SJL-*Ptprc^a^Pepc^b^*/BoyJ) mice were purchased from The Jackson Laboratory. NOD/*Scid* IL2RγC^null^ (NSG) mice were bred at City of Hope. All mouse experiments were approved by the City of Hope Institutional Animal Care and Use Committee (IACUC). The luciferase-expressing murine GL261 (GL261-Luc) and KR158B (K-Luc) glioma cells were transduced with lentivirus to produce murine IL13Rα2 (mIL13Rα2) expressing sublines (GL261-Luc-mIL13Rα2 and K-Luc-mIL13Rα2). These tumor lines were maintained in DMEM (Gibco) supplemented with 10% fetal bovine serum (Hyclone Laboratories), 25mM HEPES (Irvine Scientific, Santa Ana, CA) and 2mM L-glutamine (Lonza). Cell surface expression of mIL13Rα2 was authenticated by flow cytometry and immunofluorescence imaging.

Patient-derived glioma cells (PBT030-2-ffLuc and PBT103-2) were isolated from GBM patient resections under protocols approved by the COH IRB and maintained as described previously (*16*). All tumor lines were authenticated for the desired antigen/marker expression by flow cytometry and cells were tested for mycoplasma and maintained in culture for less than 1-3 months.

### CAR T cell Production

#### Human CAR T cells

Naïve and memory T cells were isolated from healthy donors at City of Hope under protocols approved by the COH IRB(*30, 31*). The construct of IL13Rα2-targeted CAR and CAR transduction was described in previous studies(*30, 32*),(*33*). In brief, primary T cells were stimulated with Dynabeads Human T Expander CD3/CD28 (Gibco) (T cells : beads = 1:2) for 24 hours and transduced with CAR lentivirus (multiplicity of infection [MOI] = 0.5). 7 days after CAR transduction, CD3/CD28 beads were removed and cells were resuspended and expanded in X-VIVO 15 media (Lonza) containing 10% FCS, 50 U/ml recombinant human IL-2 (Novartis), and 0.5 ng/ml recombinant human IL-15 (CellGenix) for additional 10-15 days before proceeding to cryopreservation for future experiments.

#### Murine CAR T cells

The murine IL13Rα2 CAR T cells were generated as previously reported(*13*). Briefly, T cells were isolated from spleens of naïve wild-type C57BL/6J or CD45.1 mice with EasySep Mouse T cell Isolation Kit (STEMCELL Technologies) and stimulated with Dynabead Mouse T-Activator CD3/CD28 beads (Gibco) at a 1:1 ratio. T cells were transduced on RetroNectin-coated plates (Takara Bio USA) using IL13Rα2-CAR retrovirus-containing supernatants. Cells were then expanded for 4 days in RPMI-1640 (Lonza) supplemented with 10% FBS (Hyclone Laboratories), 55 mM 2-mercaptoethanol (Gibco), 50 U/mL recombinant human-IL-2 (Novartis), and 10 ng/mL recombinant murine IL-7 (Peprotech). Before *in vitro* and *in vivo* experiments, T cells were debeaded and CAR expression was determined by flow cytometry.

TGFβR2KO-IL13Rα2-CAR T cells were isolated from spleen, stimulated with CD3/CD28 beads, and transduced with IL13Rα2-CAR retrovirus-containing supernatant as described above. After 48 hours of incubation with retroviral supernatant, CD3/CD28 beads were magnetically removed from the CAR T cells. A pool of two TGFβR2 gRNA (5’-gaagccgcaugaagucugcg-3’ and 5’-gucauggccagcgacaaugg-3’) were introduced via electroporation (Lonza), and then cells rested for 2 hours at 37°C, 5% CO_2_. CD3/CD28 beads were re-added to the culture medium, and 50% of the media was changed daily for the next 72 hours. On the 8^th^ day of production, the CAR T products were debeaded and used for subsequent experiments. For experiments comparing TGFβR2KO-IL13Rα2-CAR T cells to IL13Rα2-CAR T cells, the IL13Rα2-CAR T were electroporated without gRNA, and used at Day 8. Knockout efficiency was later determined by TIDE assay.

### qRT-PCR analysis

RNA was isolated from myelin-removed mouse brain tissue or CD3/CD28 bead-stimulated human T cells using the RNeasy Mini Kit (Qiagen). cDNA was reverse transcribed using the SuperScript VILO Mastermix (Life Technologies) according to the manufacturer’s instructions. qPCR reactions were performed as previously described(*31*). Primers are available in Supplementary Table 1.

### In vivo studies

All mouse experiments were performed using protocols approved by the City of Hope IACUC. Orthotopic GBM models were generated as previously described(*34*). Orthotopic tumor model was established by stereotactically implanting 1×10^5^ tumor cells intracranially (i.c.) into the right forebrain of 8–10 week-old C57BL/6J mice. Engraftment was verified by bioluminescent imaging one day prior to treatments.

#### CRISPR/Cas TGFβR2KO IL13Rα2 CAR T therapy studies

14 days after tumor injection, mice were treated intracranially with 1x10^6^ IL13Rα2 or TGFβR2KO IL13Rα2 CAR T cells.

#### TGFβR1 inhibitor plus IL13Rα2 CAR T therapy studies

four days after tumor implantation, mice were given oral dose of TGFβR1 inhibitor (LY3200882, 210 mg/kg) twice daily for 26 days. On day 14, only the morning dose of TGFβR1 inhibitor was administered, followed by IL13Rα2 CAR T cells (1x10^6^) injection to the intracranial tumor bed.

Tumor burden was monitored with SPECTRAL LagoX (Spectral Instruments Imaging) and analyzed using Aura software (v2.3.1, Spectral Instruments Imaging). Survival curves were generated by GraphPad Prism Software (v8). Weights were measured before and during treatment for all experiments as read out for any potential toxicity.

Mice were also monitored by the Center for Comparative Medicine at City of Hope for survival and any symptoms related to tumor progression, with euthanasia applied according to the American Veterinary Medical Association Guidelines.

For rechallenge experiments, clearance of tumor was verified by bioluminescent imaging prior to tumor rechallenge, where mice were injected with 10^4^ K-Luc or 5x10^4^ GL261-Luc cells.

### In vitro stimulation and cytotoxicity assays

#### For in vitro stimulation assays

human CAR T cells from healthy donors were thawed and stimulated with CD3/CD28 beads (Invitrogen) at a ratio of 1:1, in the presence or absence of 10 ng/mL recombinant TGFβ1 or 3 µM TGFβR1i (LY3200882, Lilly). At the end of four days, cells were either lysed for protein (western blot) or RNA (qPCR), or stained with fluorescent antibodies (PD1, CD8) for flow cytometry.

#### For in vitro rechallenge assays

human CAR T cells from healthy donors or patients were thawed and cocultured with low or high IL13Ra2+ tumors as described previously(*35*). After 7 days (high antigen) or 9 days (low antigen), remaining tumors and T cells were stained with CD3, CD4, and CD19 antibodies, then counted by flow cytometry. For flow sorting, cells were separated by flow cytometry into CD3+CD19+CD4+ or CD3+CD19+CD8+ populations and subsequently lysed for downstream RNA analysis by Nanostring.

#### For in vitro killing with TGFβR2KO-IL13Rα2-CAR T

1500 T cells were plated with 6000 IL13Rα2+ tumor cells for 24 hours. Viable tumor and T cells were stained with CD3, CD4, CD8, CD19 antibodies and then counted by flow cytometry.

All samples were acquired on MACSQuant10 Analyzer (Miltenyi Biotec) and analyzed with FlowJo software (v10.7) and GraphPad Prism (v8).

### Patient ELISA assays

Tumor cyst fluid (TCF) and cerebrospinal fluid (CSF) were collected as previously described (*15*). Samples were analyzed using a precoated ELISA for human TGFβ1 (R&D Systems) according to the manufacturer’s protocol. TCF and CSF samples were assayed without dilution. Each batch of samples was analyzed using a four-parameter logistic regression generated from that batch’s standard curve.

### TME characterization

Brains from euthanized mice were removed at the indicated time-points, and a rodent brain matrix was used to cut along the coronal and sagittal planes to obtain a 4x4 mm section, centered around the injection site. These sections were minced manually, then passed through a 40 µm filter. Myelin was removed using Myelin Removal Beads II and LS magnetic columns (Miltenyi Biotec) according to the manufacturer’s instructions, then cells were counted. Cells were stained with fluorescently labelled CD3, CD69, CD8, CD4, CD19, CXCR3, CD45.1, CD11b, CD86, MHCII, MHCI, Ki67, or IFNγ. All samples were acquired on MACSQuant Analyzer10 (Miltenyi Biotec) and analyzed with FlowJo software (v10.7) and GraphPad Prism (v8).

For gene expression analysis of TME, the remaining cells were lysed for RNA.

### NanoString gene expression analysis

RNA was purified from flow-sorted CD4^+^ or CD8^+^ sorted cells using the RNEasy Mini kit, following the manufacturer’s instructions (Qiagen, Germantown, MD, USA). RNA samples were subsequently quantified and qualified using Nanodrop 1000 spectrophotometer (ThermoFisher, Waltham, MA, USA). The subsequent Nanostring analysis was performed at concentrations of 35ng/well and 25ng/well respectively for CD3^+^ cells and CD11b^+^ cells.

Samples were analyzed based on the nCounter^®^ human CAR-T Characterization gene expression panel (NanoString Technologies, Seattle, WA, USA): Hybridation reaction was performed for 18h at 65°C. Fully automated nCounter FLEX analysis system; composed of an automated nCounter^®^ Prep station and the nCounter^®^ Digital Analyzer optical scanner (NanoString Technologies, Seattle, WA, USA) was used. Normalization was performed by using the geometric mean of the positive control counts as well as normalization genes present in the CodeSet Content: samples with normalization factors outside of the 0.3–3.0 range were excluded, samples with reference factors outside the 0.10–10.0 range were excluded as well. Gene expression analysis was performed using the nSolver v3.0 and Advanced analysis module softwares. (NanoString Technologies, Seattle, WA, USA).

The analysis on Gene ontology (GO), Kyoto encyclopedia of genes and genomes pathway, and Immunologic signatures collection (ImmuneSigDB) was performed with the full gene expression matrix on CD4 or CD8 cells separately, using GSEA function implemented in clusterProfiler package.

### Bulk RNA sequencing

Cells from dissociated mouse brains that were not used for flow cytometry characterization were pelleted and frozen. Pellets were thawed into RLT buffer, and RNA was extracted from these pellets using the RNEasy Mini kit according to the manufacturer’s instructions.

RNA-seq library preparation was conducted as previously described(*12*). RNA-Seq reads were trimmed to remove sequencing adapters using Trimmomatic (*36*)and polyA tails using FASTP. (*37*). The processed reads were mapped back to the mouse genome (mm10) using STAR (*38*) software (v. 2.6.0.a). The HTSeq (*39*)software (v.0.11.1) was applied to generate the count matrix, with default parameters. Differential expression analysis was conducted by adjusting read counts to normalized expression values using TMM normalization method in edgeR (*40, 41*). Briefly, for between-treatment comparison, general linear models were applied to identify DEGs using TMM normalization expression level as the dependent variable, and treatment as the independent variable. Genes with an FDR-adjusted p-value less than 0.05 and with a fold change (FC) greater than 2 or less than 0.5 were considered as significant up- and down-regulated genes, respectively. Pathway analysis was conducted using GSEA algorithm implemented in clusterProfiler (*42–45*) (v.4.14.3) package in R where a ranked list of whole genes according to their log2 fold change and p-values are provided.

### Single cell RNA-sequencing

Patients were segregated into a high TGFβ (n = 11) or low TGFβ (n = 6) cohort based on the ELISA for TGFβ at Cycle 1 Day 0. For both the CAR product and the tumor microenvironment analysis, the datasets were trimmed to only include patients with matching TGFβ ELISA data from the TCF compartment at Cycle 1 Day 0. For CAR T product analyses scRNAseq data sets were obtained from GEO (GSE255850, (*15*)). For the tumor microenvironment analysis, previously identified immune clusters (*11*) were subsetted from tumor cells using Seurat (v5.1.0) and reclustered separately, either as an immune or a tumor Seurat object. Pathway analysis was conducted using the escape package (v2.1.4) in R (v4.4.1).

### TCGA Glioblastoma Multiforme public cohort mining

The public cohort mining of TGFB1 mRNA seq gene expression comparing tumor vs normal referenced in Supplemental Figure 1A from QIAGEN OmicSoft OncoLand database TCGA_B38_GC36 were used under license for this study.

### Statistical analysis

Statistical significance was determined using Student *t*-test (two groups) or one-way ANOVA analysis with a Bonferroni correction (three or more groups), unless noted otherwise. The significance level for figure 1D was calculated using a linear regression between groups, to allow each patient to have a base level and thereby accounting for the repeated measures from each patient. Survival was plotted using a Kaplan-Meier survival curve and statistical significance was determined by the Log-rank (Mantel-Cox) test. All analyses were carried out either using GraphPad Prism software (v5) or R. In particular distribution curves were estimated using the base R density function. For comparisons showing individual measures between groups, data are presented as with horizontal lines highlighting means and vertical lines ± SEM. Statistical significance of relevant comparisons is indicated by: *, *p* < 0.05; **, *p* < 0.01; ***, *p* < 0.001; ****, *p* <0.0001.

## List of Supplementary Materials

Supplementary Methods

Fig. S1. TGFβ pathway in GBM patients.

Fig. S2. Inhibition of TGFβ signaling enhances the CAR T function and proliferation against low antigen expressing tumors.

Fig. S3: Effect of CAR T cell treatment in large established murine gliomas.

Fig. S4: Changes in weight in mice treated with combination CAR T and TGFβR1i.

Fig. S5: Blocking TGFβ signaling enhances CAR T antitumor function in vivo.

Fig. S6: Combination TGFβ blockade and CAR T cells induce long-term immunity.

Fig. S7: Expression of activated cytotoxic T cells in tumor environment.

Fig. S8: Characterization of TGFβR2KO-IL13Rα2 T cells.

Supplementary Table 1: Primers used for qPCR and TIDE assay.

## Acknowledgments

The authors would like to thank Kyla Elizabeth Driscoll and Eli Lilly and Company for providing the TGFβR1 inhibitor LY3200882 and Vivian Chiu for their technical assistance.

## Funding

This work was supported in part by research grants from the

National Institutes of Health R01CA220693 (CEB)

Catherine Ivy Foundation (CEB/DA)

Mann Foundation (CEB)

B*Cured (DA)

Norris Foundation (DA)

Cancer Center Support Grant P30CA33572

Mustang Bio., Inc. (CEB)

## Author contributions

DA, RAW, and CEB contributed to overall project design and wrote the manuscript. DA, RAW and CEB contributed to methodology development. DA, RAW, MSM, DB, RS, JRP, and JDJ contributed to acquisition of data. JCH contributed to data acquisition and biostatistics. DA, RAW, and CEB did analysis and interpretation of data. YY contributed to public data mining and acquisition. MSM, WC, ZW contributed to studies related to CRISPR/Cas. HMN, MC, and XW supervised bioinformatics analysis. NEB, BB, SJF assisted in reviewing the manuscript. DA, RAW and CEB supervised the overall study.

## Competing interests

Patents associated with IL13Rα2-CAR T have been licensed by Mustang Bio., Inc., for which S.J.F. and C.E.B. receive royalty payments. All other authors declare no potential conflicts of interest.

## Data and materials availability

All data associated with this study are present in the paper or the Supplementary Materials. All datasets related to scRNA-seq and bulk RNA-seq data will be publicly accessible upon publication.

## References and Notes

1. S. J. Bagley et al., Intrathecal bivalent CAR T cells targeting EGFR and IL13Ralpha2 in recurrent glioblastoma: phase 1 trial interim results. Nat Med 30, 1320–1329 (2024).

2. S. J. Bagley et al., Intracerebroventricular bivalent CAR T cells targeting EGFR and IL-13Ralpha2 in recurrent glioblastoma: a phase 1 trial. Nat Med 31, 2778–2787 (2025).

3. C. E. Brown et al., Locoregional delivery of IL-13Rα2-targeting CAR-T cells in recurrent high-grade glioma: a phase 1 trial. Nature Medicine 30, 1001–1012 (2024).

4. J. Han, C. A. Alvarez-Breckenridge, Q. E. Wang, J. Yu, TGF-beta signaling and its targeting for glioma treatment. Am J Cancer Res 5, 945–955 (2015).

5. S. Mariathasan et al., TGFbeta attenuates tumour response to PD-L1 blockade by contributing to exclusion of T cells. Nature 554, 544–548 (2018).

6. B. V. Park et al., TGFbeta1-Mediated SMAD3 Enhances PD-1 Expression on Antigen-Specific T Cells in Cancer. Cancer Discov 6, 1366–1381 (2016).

7. E. Batlle, J. Massague, Transforming Growth Factor-beta Signaling in Immunity and Cancer. Immunity 50, 924–940 (2019).

8. D. Akhavan et al., CAR T cells for brain tumors: Lessons learned and road ahead. Immunol Rev 290, 60–84 (2019).

9. L. Qiang et al., Transforming Growth Factor-beta Blockade in Pancreatic Cancer Enhances Sensitivity to Combination Chemotherapy. Gastroenterology 165, 874–890 e810 (2023).

10. C. Flood et al., Transforming growth factor-beta1 in the cerebrospinal fluid of patients with subarachnoid hemorrhage: titers derived from exogenous and endogenous sources. J Cereb Blood Flow Metab 21, 157–162 (2001).

11. S. Gholamin et al., Overcoming myeloid-driven resistance to CAR T therapy by targeting SPP1. bioRxiv, (2025).

12. D. Alizadeh et al., IL15 Enhances CAR-T Cell Antitumor Activity by Reducing mTORC1 Activity and Preserving Their Stem Cell Memory Phenotype. Cancer Immunol Res 7, 759–772 (2019).

13. D. Alizadeh et al., IFNγ Is Critical for CAR T Cell-Mediated Myeloid Activation and Induction of Endogenous Immunity. Cancer Discov 11, 2248–2265 (2021).

14. A. J. Gunderson et al., TGFbeta suppresses CD8(+) T cell expression of CXCR3 and tumor trafficking. Nat Commun 11, 1749 (2020).

15. C. Brown, et al., Phase I Trial Evaluating Locoregionally-delivered IL13Rα2-targeting CAR T Cells in High-Grade Glioma. (2023).

16. C. E. Brown et al., Glioma IL13Ralpha2 is associated with mesenchymal signature gene expression and poor patient prognosis. PloS one 8, e77769 (2013).

17. Q. C. Qiu et al., CHI3L1 promotes tumor progression by activating TGF-beta signaling pathway in hepatocellular carcinoma. Sci Rep 8, 15029 (2018).

18. T. A. Yap et al., First-In-Human Phase I Study of a Next-Generation, Oral, TGFbeta Receptor 1 Inhibitor, LY3200882, in Patients with Advanced Cancer. Clin Cancer Res 27, 6666–6676 (2021).

19. E. Malek et al., The TGFbeta type I receptor kinase inhibitor vactosertib in combination with pomalidomide in relapsed/refractory multiple myeloma: a phase 1b trial. Nat Commun 15, 7388 (2024).

20. J. H. Ahn et al., Clinical Activity of TGF-beta Inhibitor Vactosertib in Combination with Imatinib in Desmoid Tumors: A Multicenter Phase Ib/II Study. Clin Cancer Res 30, 1457–1465 (2024).

21. A. Maier, A. L. Peille, V. Vuaroqueaux, M. Lahn, Anti-tumor activity of the TGF-beta receptor kinase inhibitor galunisertib (LY2157299 monohydrate) in patient-derived tumor xenografts. Cellular oncology 38, 131–144 (2015).

22. T. A. Yap et al., PF-06952229, a selective TGF-beta-R1 inhibitor: preclinical development and a first-in-human, phase I, dose-escalation study in advanced solid tumors. ESMO Open 9, 103653 (2024).

23. N. Tang, et al., TGF-beta inhibition via CRISPR promotes the long-term efficacy of CAR T cells against solid tumors. JCI Insight 5, (2020).

24. A. Gieryng, D. Pszczolkowska, K. A. Walentynowicz, W. D. Rajan, B. Kaminska, Immune microenvironment of gliomas. Laboratory investigation; a journal of technical methods and pathology 97, 498–518 (2017).

25. C. C. Kloss et al., Dominant-Negative TGF-beta Receptor Enhances PSMA-Targeted Human CAR T Cell Proliferation And Augments Prostate Cancer Eradication. Mol Ther 26, 1855–1866 (2018).

26. V. Narayan et al., PSMA-targeting TGFbeta-insensitive armored CAR T cells in metastatic castration-resistant prostate cancer: a phase 1 trial. Nat Med 28, 724–734 (2022).

27. K. Li et al., Dominant-negative transforming growth factor-beta receptor-armoured mesothelin-targeted chimeric antigen receptor T cells slow tumour growth in a mouse model of ovarian cancer. Cancer Immunol Immunother 72, 917–928 (2023).

28. N. Li et al., Armored bicistronic CAR T cells with dominant-negative TGF-beta receptor II to overcome resistance in glioblastoma. Mol Ther 32, 3522–3538 (2024).

29. A. J. Hou, Z. L. Chang, M. H. Lorenzini, E. Zah, Y. Y. Chen, TGF-beta-responsive CAR-T cells promote anti-tumor immune function. Bioeng Transl Med 3, 75–86 (2018).

30. C. E. Brown et al., Optimization of IL13Ralpha2-Targeted Chimeric Antigen Receptor T Cells for Improved Anti-tumor Efficacy against Glioblastoma. Mol Ther 26, 31–44 (2018).

31. D. Wang, et al., Glioblastoma-targeted CD4+ CAR T cells mediate superior antitumor activity. JCI insight 3, e99048 (2018).

32. C. E. Brown et al., Regression of Glioblastoma after Chimeric Antigen Receptor T-Cell Therapy. The New England journal of medicine 375, 2561–2569 (2016).

33. D. Wang et al., Chlorotoxin-directed CAR T cells for specific and effective targeting of glioblastoma. Sci Transl Med 12, (2020).

34. C. E. Brown et al., Recognition and killing of brain tumor stem-like initiating cells by CD8+ cytolytic T cells. Cancer Res 69, 8886–8893 (2009).

35. D. Wang et al., In Vitro Tumor Cell Rechallenge For Predictive Evaluation of Chimeric Antigen Receptor T Cell Antitumor Function. J Vis Exp, (2019).

36. A. M. Bolger, M. Lohse, B. Usadel, Trimmomatic: a flexible trimmer for Illumina sequence data. Bioinformatics 30, 2114–2120 (2014).

37. S. Chen, Y. Zhou, Y. Chen, J. Gu, fastp: an ultra-fast all-in-one FASTQ preprocessor. Bioinformatics 34, i884–i890 (2018).

38. A. Dobin et al., STAR: ultrafast universal RNA-seq aligner. Bioinformatics 29, 15–21 (2013).

39. S. Anders, W. Huber, Differential expression analysis for sequence count data. Genome Biol 11, R106 (2010).

40. M. D. Robinson, D. J. McCarthy, G. K. Smyth, edgeR: a Bioconductor package for differential expression analysis of digital gene expression data. Bioinformatics 26, 139–140 (2010).

41. D. J. McCarthy, Y. Chen, G. K. Smyth, Differential expression analysis of multifactor RNA-Seq experiments with respect to biological variation. Nucleic Acids Res 40, 4288–4297 (2012).

42. G. Yu, L. G. Wang, Y. Han, Q. Y. He, clusterProfiler: an R package for comparing biological themes among gene clusters. OMICS 16, 284–287 (2012).

43. T. Wu et al., clusterProfiler 4.0: A universal enrichment tool for interpreting omics data. Innovation (Camb) 2, 100141 (2021).

44. S. Xu et al., Using clusterProfiler to characterize multiomics data. Nat Protoc 19, 3292–3320 (2024).

45. G. Yu, Thirteen years of clusterProfiler. Innovation (Camb) 5, 100722 (2024).

