## Supplemental Data for "Therapeutic Targeting of the TGFβ Pathway Augments CAR T cell Efficacy in Glioblastoma"

### **Supplementary Methods**

#### ***Patient Immunohistochemistry***

4 μm sections of either normal brain or GBM tumor were obtained from City of Hope's Tumor Biobank. Sequential slides were stained with H&E and pSmad2 antibody. Slides of pSmad2 stains were scanned with a Hamamatsu slide scanner, and then positively stained nuclei were counted using VisioPharm software. H&E stains were examined by the City of Hope pathologist to identify tumor areas. Tumor areas were then manually scrapped off the slide, and RNA was extracted from these curls using the RNEasy FFPE Kit according to the manufacturer's instructions. RNA sequencing was performed as described above.

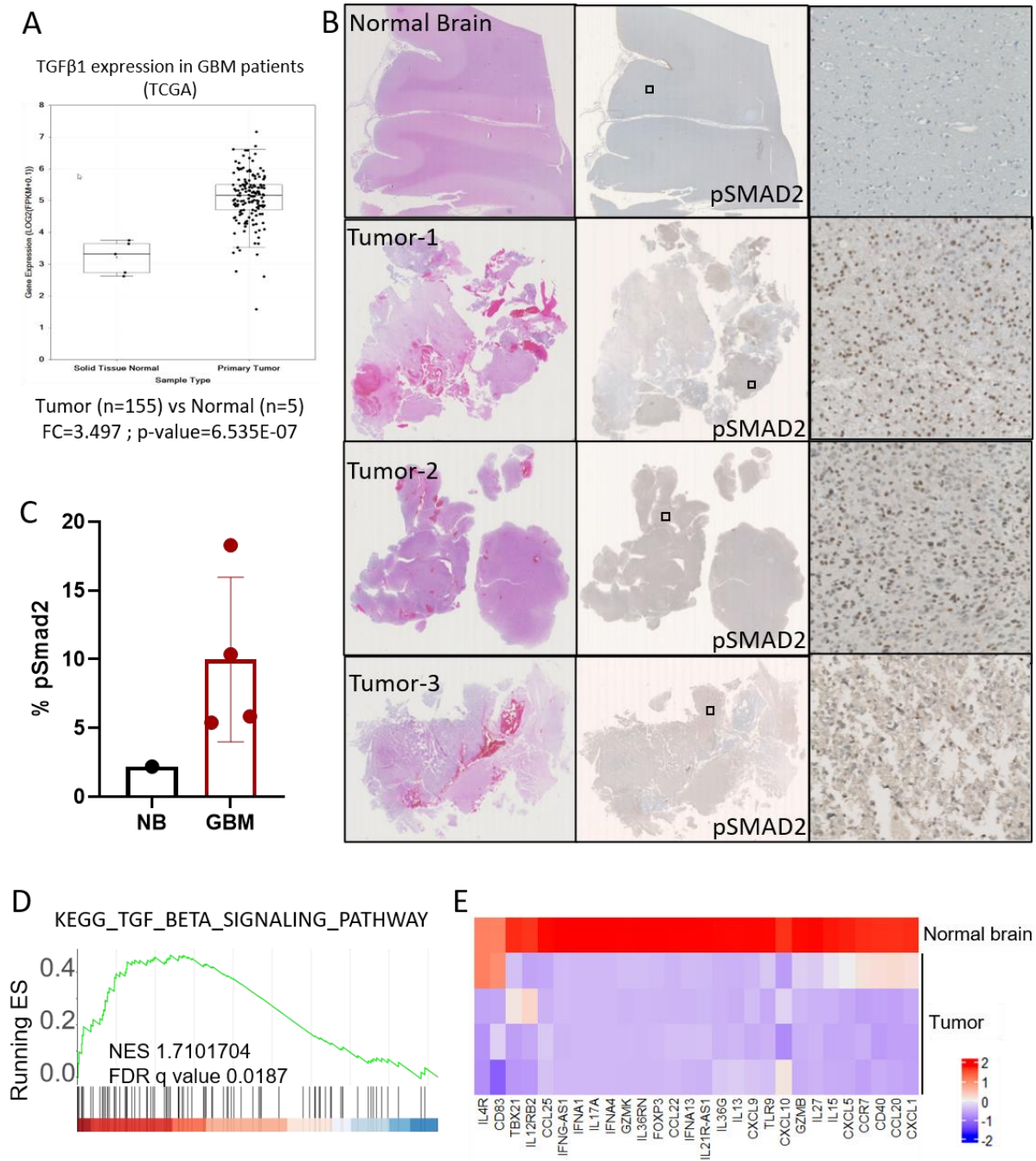

**Fig. S1. TGFβ pathway in GBM patients.** (A) TCGA database shows *TGFβ1* expression levels in normal and GBM tissues. (B) Histology of normal brain and tumor tissues using H&E and IHC staining of pSMAD2. Width of inset box and enlargement is 500 μm. (C) Graph shows quantification of pSMAD2 nuclei over total nuclei. (D) TGFβ pathway enrichment analysis in

GBM tumor tissues. (E) Heatmap of normalized expression of selected genes on patient tumor samples (n=4) vs normal brain.

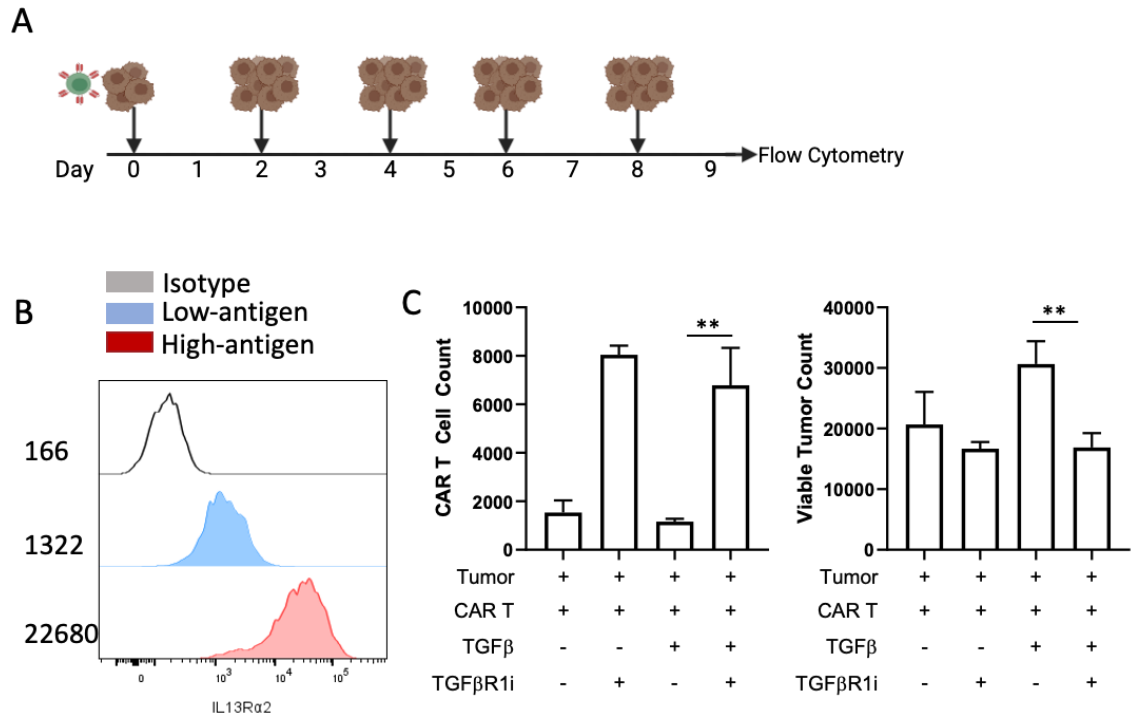

**Fig. S2. Inhibition of TGFβ signaling enhances the CAR T function and proliferation against low antigen expressing tumors. (A)** Schematic of experimental design. **(B)** Histograms demonstrate IL13Rα2 expression levels by flow cytometry in tumor cells lines. Numbers indicate MFI. **(C)** Graphs show CAR T cell expansion (left) and viable tumor counts (right) after multiple tumor rechallenge in the presence or absence of TGFβ (10ng/ml) or TGFβR1 inhibitor (3 μM), against a patient derived glioma tumor expressing low levels of IL13Rα2.

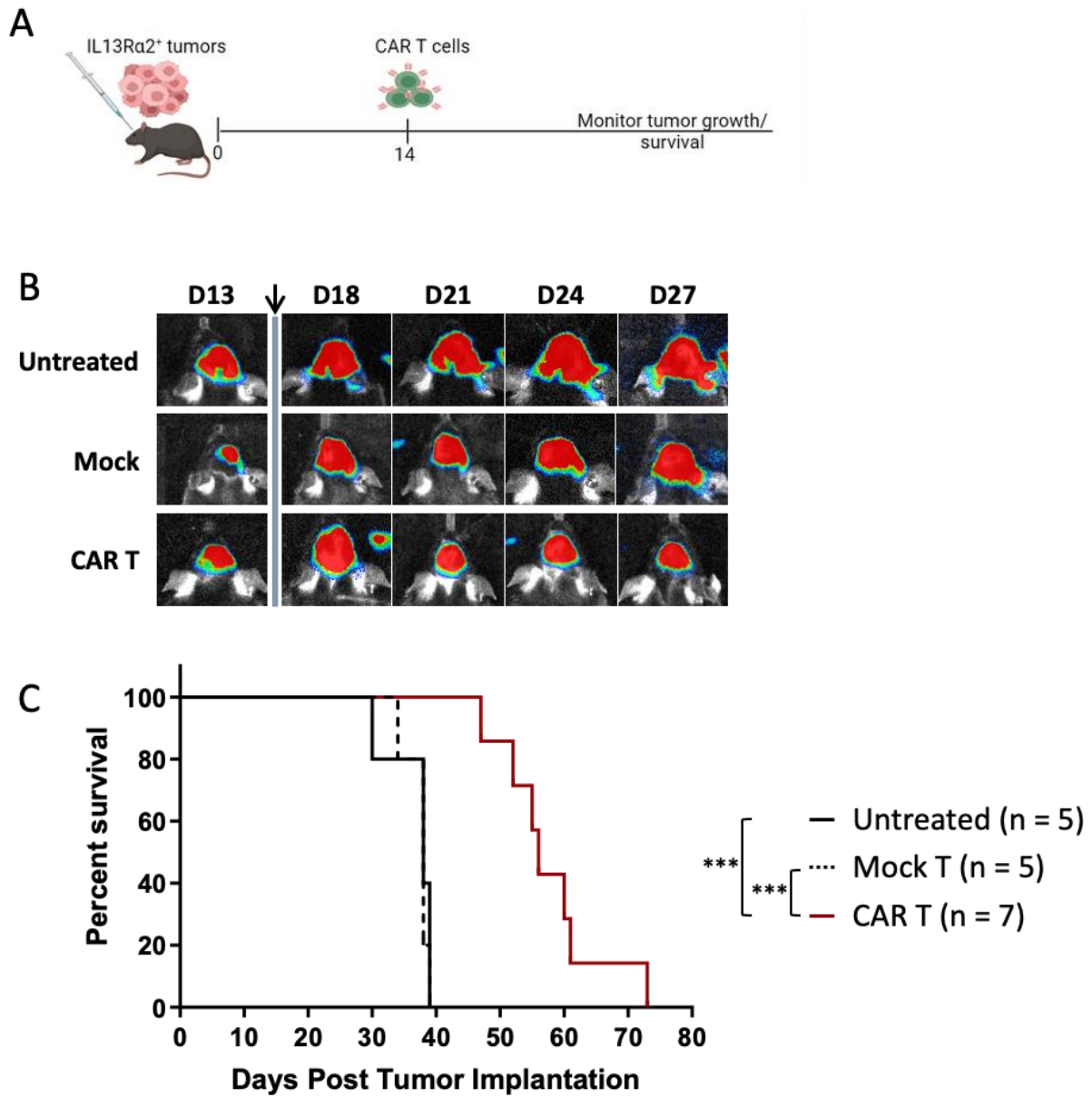

**Fig. S3: Effect of CAR T cell treatment in large established murine gliomas. (A)** Schematic of experimental design. **(B)** Representative BLI images of tumor progression after mock or CAR T treatment delivered intratumorally. **(C)** Kaplan Meier graph shows overall survival of mice in untreated, mock, or CAR T treated ($1 \times 10^6$ cells) groups.

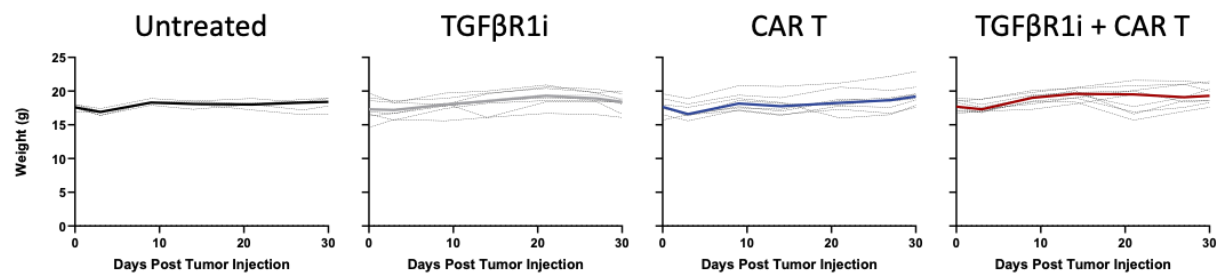

**Fig. S4: Changes in weight in mice treated with combination CAR T and TGFβR1i.** Graphs show changes in body weight measured over time.

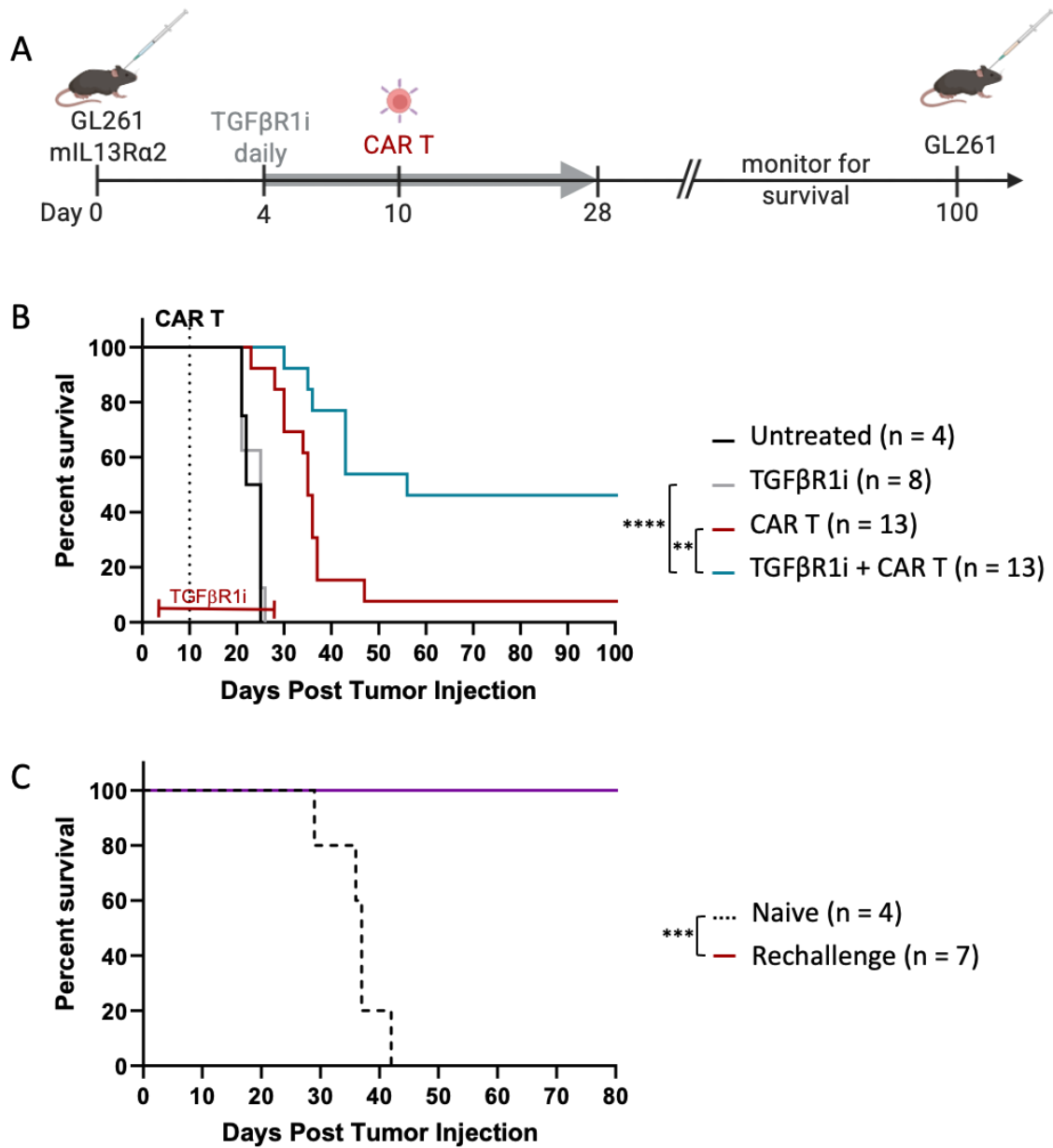

**Fig. S5: Blocking TGFβ signaling enhances CAR T antitumor function in vivo.**

(A) Schematic of experimental design. Mice bearing glioma tumors (10-day old GL261-mIL13Rα2+) were treated with TGFβR1 inhibitor daily before and after IL13Rα2-CAR T treatment. (B) Kaplan Meier graph shows overall survival of mice in untreated, TGFβR1i alone, CAR T ($1 \times 10^6$ cells) alone, or combination TGFβR1i plus CAR T treated groups in GL261

intracranial tumor model. (C) Kaplan Meier graph shows rechallenge of tumor-free survivors from B with antigen-negative GL261.

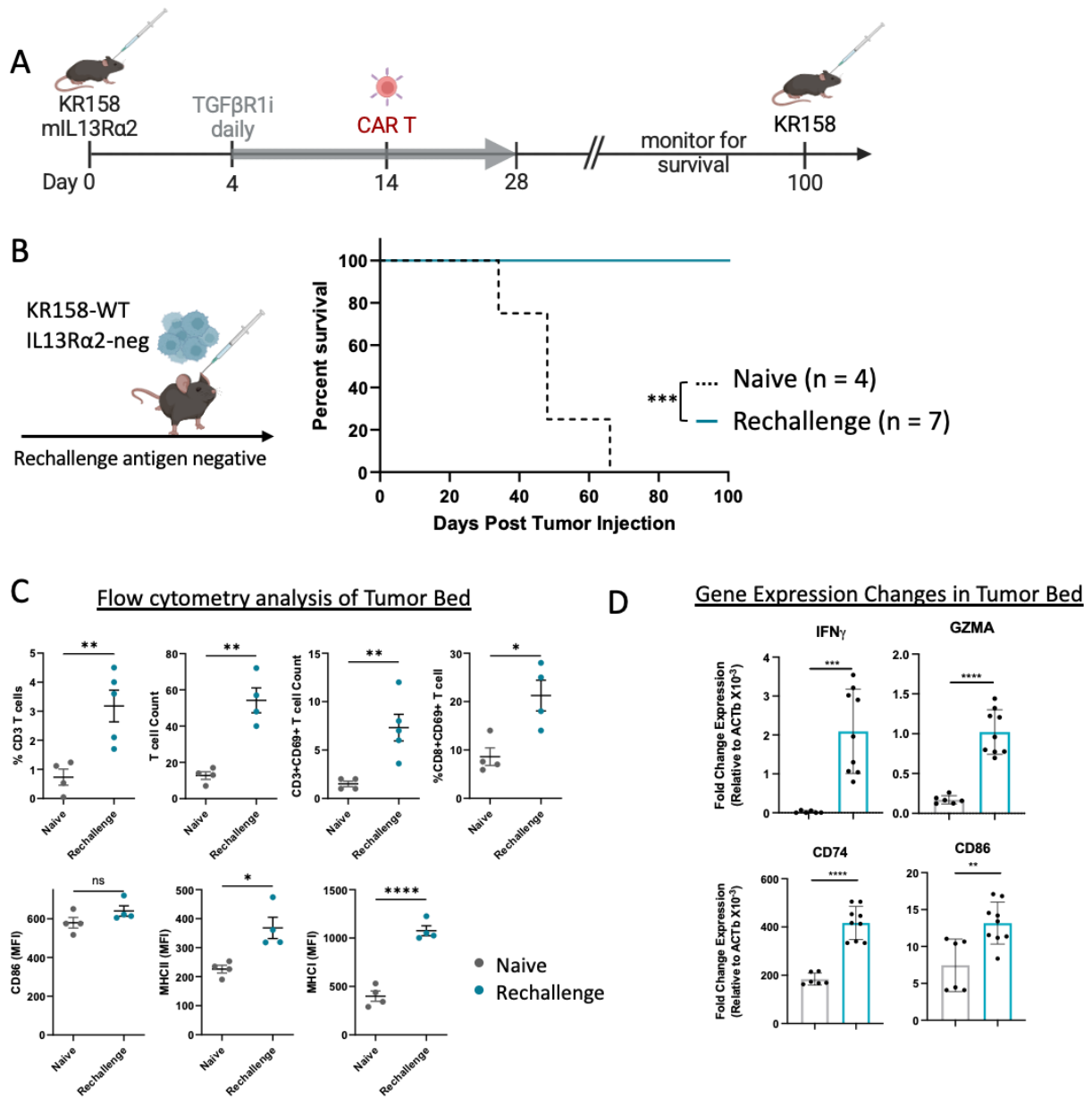

**Fig. S6: Combination TGFβ blockade and CAR T cells induce long-term immunity. A.** Schematic of experimental design. **B.** Kaplan Meier survival curve of mice challenged with IL13Rα2 antigen negative KR158 tumors. **C.** Graphs show change in immune landscape in the TME two days after tumor challenge. **D.** qPCR analyses of total TME shows changes in expression of genes associated with immune activation.

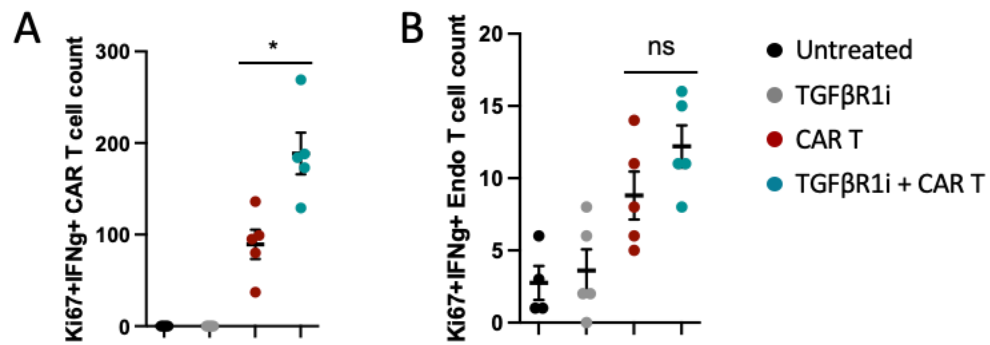

**Fig. S7: Expression of activated cytotoxic T cells in tumor environment.** Graphs quantify flow cytometry analysis of cytotoxic (IFNg+Ki67+) **A.** CAR T cell and **B.** endogenous T cells in each treatment groups.

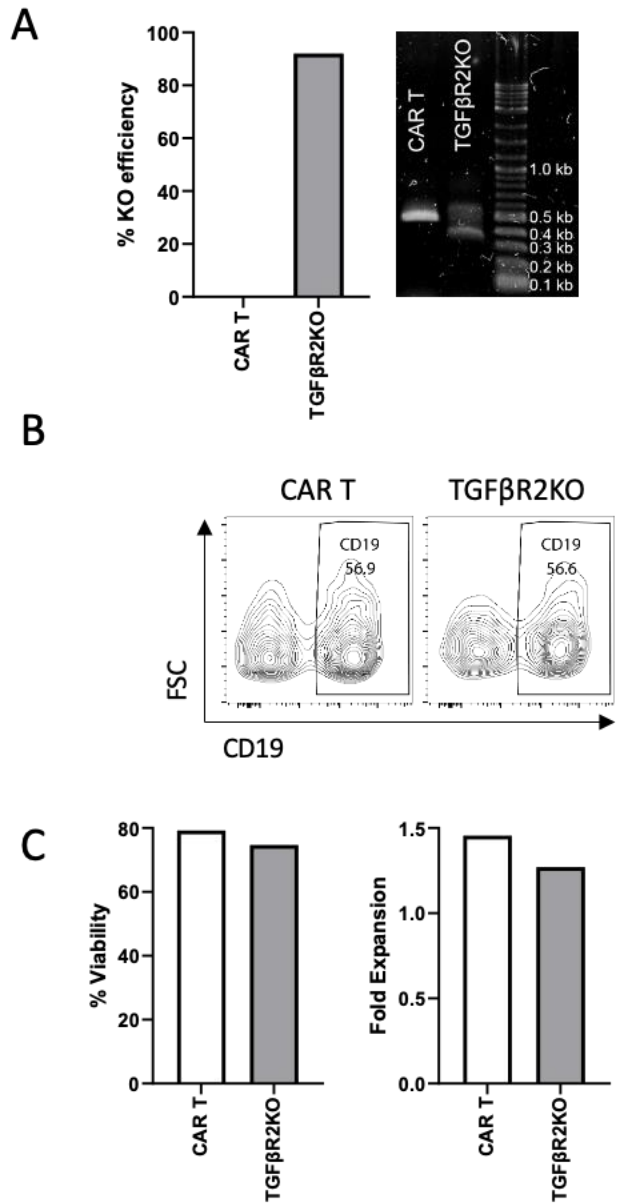

**Fig. S8: Characterization of TGFβR2KO-IL13Rα2 T cells.** **A.** Graph demonstrates knockout efficiency using TIDE assay and agarose gel. **B.** Flow cytometry analysis shows transduction efficiency of the CAR construct in TGFβR2KO or standard-IL13Rα2-CAR T cells. **C.** Graphs compare viability (left) and fold expansion (right) in TGFβR2KO or standard-IL13Rα2-CAR T cells.

**Supplementary Table 1: Primers used for qPCR and TIDE assay.**

| Gene | Species | Forward | Reverse |
| --- | --- | --- | --- |
| *GAPDH* | human | CTCATGACCACAGTCCATGCC | AGGCCATGCCAGTGAGCTTC |
| *PDCDI* | human | AAACCCTGGTGGTTGGTGTC | CTCCTATTGTCCCTCGTGCG |
| *PRF1* | human | CTGGGTGGAGGCGTTGAAG | GGCTGGACGTGACTCCTAAG |
| *Actb* | mouse | AGCTCATTGTAGAAGGTGTGG | GTGGGAATGGGTCAGAAGG |
| *Cd74* | mouse | AGATGCGGATGGCTACTCC | TCATGTTGCCGTACTTGGTAAC |
| *Cd86* | mouse | TCAATGGGACTGCATATCTGCC | GCCAAAATACTACCAGCTCACT |
| *Gzma* | mouse | TGCTGCCCACTGTAACGTG | GGTAGGTGAAGGATAGCCACAT |
| *Ifng* | mouse | ACAGCAAGGCGAAAAAGGATG | TGGTGGACCACTCGGATGA |
| *Tgfbr2* | mouse | TTGATGTCCAGCTGCTCCAG | ATGCCATGCCATTTGTGCAG |

| Gene | Species | Taqman Assay ID |
| --- | --- | --- |
| *GAPDH* | human | Hs02758991_g1 |
| *GZMA* | human | Hs00989184_m1 |
| *IFNG* | human | Hs00989291_m1 |
